# Hidden dynamic signatures drive substrate selectivity in the disordered phosphoproteome

**DOI:** 10.1101/866558

**Authors:** Min-Hyung Cho, James O. Wrabl, James Taylor, Vincent J. Hilser

## Abstract

Phosphorylation sites are hyper-abundant in the disordered proteins of eukaryotes, suggesting that conformational dynamics (or heterogeneity) may play a major role in determining to what extent a kinase interacts with a particular substrate. In biophysical terms, substrate selectivity may be determined not just by the structural and chemical complementarity between the kinase and its protein substrates, but also by the free energy difference between the conformational ensembles that are recognized by the kinase and those that are not. To test this hypothesis, we developed an informatics framework based on statistical thermodynamics, which allows us to probe for dynamic contributions to phosphorylation, as evaluated by the ability to predict Ser/Thr/ Tyr phosphorylation sites in the disordered proteome. Essential to this framework is a decomposition of substrate sequence information into two types: vertical information encoding conserved kinase specificity motifs and horizontal (distributed) information encoding substrate conformational dynamics that are embedded, but often not apparent, within position specific conservation patterns. We find not only that conformational dynamics play a major role, but that they are the dominant contribution to substrate selectivity. In fact, the main substrate classifier distinguishing selectivity is the magnitude of change in compaction of the disordered chain upon phosphorylation. Thus, in addition to providing fundamental insights into the underlying mechanistic consequences of phosphorylation across the entire proteome, our approach provides a novel statistical thermodynamic strategy for partitioning any sequence-based search into contributions from direct chemical and structural complementarity and those from changes in conformational dynamics. Using this framework, we developed a high-performance open-source phosphorylation site predictor, PHOSforUS, which is freely available at https://github.com/bxlab/PHOSforUS.

## 1. Introduction

Phosphorylation is the most common post-translational modification in eukaryotic proteomes (1, 2), and has been demonstrated to mediate key biological functions, including signaling (3), nutrient sensing (4), and protein conformational change (5). In spite of the universal recognition of its importance, a significant gap in our knowledge has prevented a general mechanistic understanding of how phosphorylation mediates these processes. Specifically, many phosphorylation sites are contained within intrinsically disordered regions (IDRs) of proteins, which due to their high sequence divergence, make it a challenge to identify phosphorylatable sites based on sequence comparisons with known sites. This knowledge gap is exacerbated by the fact that phosphorylation is both transient and reversible, producing a surprisingly low degree of consensus (6) between experimentally determined phosphorylation sites in several major databases: PhosphoELM (7), UniProt (8), and PhosphoSitePlus (9), resulting, understandably, in a concomitant degree of disagreement between phosphorylation site predictors (10–16) developed from these databases (6, 17).

Attempts to address this knowledge gap have typically involved the development of heuristics to augment the limited amount of experimentally annotated sequence sites. For example, the myriad substrates of cyclin-dependent protein kinases only appear to share a single Proline (Pro) residue immediately C-terminal to the phosphorylated site (18). However, it was recognized early on that certain hydrophobic, acidic, or basic amino acid patterns were often found in the sequence neighborhood of a phosphorylation site (1, 10, 11). As a result, position specific weight matrices were developed to identify motifs predictive of kinase-specific sites, achieving a moderate degree of success when leveraged with neural network algorithms (6, 19). However, the consensus pattern approach produced significant variability, precluding practical prediction tools (6, 19).

Seminal work by Dunker and colleagues (11) revealed that phosphorylation correlates with surrounding intrinsic disorder, and explicit consideration of disorder resulted in an improved phosphorylation site predictor. Similarly, such a conformational energetic contribution was demonstrated by Elam, et al. (20) to also involve conserved polyproline II (PII) propensity of the sequence elements surrounding the phosphorylation site. Both of these observations are suggestive of a distinct role for the conformational equilibrium of the potential substrate, not only in determining the overall function of the phosphorylated protein, but also possibly in determining kinase specificity.

To test this hypothesis, we have developed a statistical thermodynamic framework that considers contributions to kinase selectivity driven either by direct recognition of sequence elements that are conserved at a particular sequence position (which we term “vertical information”) or by ensemble-averaged properties that are conserved along a sequence stretch (which we term “horizontal information”). Accounting explicitly for both types of information, “vertical” and “horizontal”, results in a predictor that exceeds performance relative to existing phosphorylation prediction methods. Indeed, our results show that the ensemble-averaged properties— equilibrium fluctuations that are encoded in horizontal information — dominate the contribution.

Furthermore, our results indicate that the sequence neighborhoods of many Serine (Ser) and Threonine (Thr) phosphorylation sites, specifically those containing Pro immediately C-terminal to the phosphorylated site (i.e. the +1 Pro sequence motif), are “energetically poised” to undergo a phosphorylation-induced change in the dimensions of the disordered ensemble, suggestive of a direct link between the conformational dimensions of the disordered substrate and its ability to be recognized and phosphorylated.

## 2. Results

### 2.1. Phosphorylation equilibria can be reflected by two types of sequence information

Enrichment of disorder around phosphorylation sites has been noted previously (11), suggesting the necessity for widespread coupled folding-binding of the disordered substrate in order to become phosphorylated (Fig. 1, top). If this is the case, it would be desirable to develop a strategy that accounts for the free energy change associated with narrowing or expanding the conformational ensemble (Fig. 1, blue box). This would involve selecting, from among the entire conformational ensemble, the sub-ensemble wherein the residues that are recognized by the kinase are in the proper orientation for kinase recognition. For that sub-ensemble, recognition and binding would then be based on classic notions of shape and chemical complementarity (Fig. 1, red box). Thus, the recognition of conformationally heterogeneous substrates by kinases can be viewed as being due to two distinct physical processes: a contribution arising from the energy difference between the substrate sub-ensemble that can be phosphorylated and the sub-ensemble that cannot, and a contribution from the intrinsic ability of the kinase to recognize the substrate, a scenario that is captured by the equilibrium:

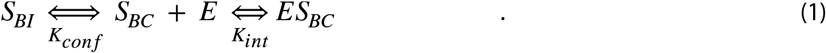

**Figure 1.**
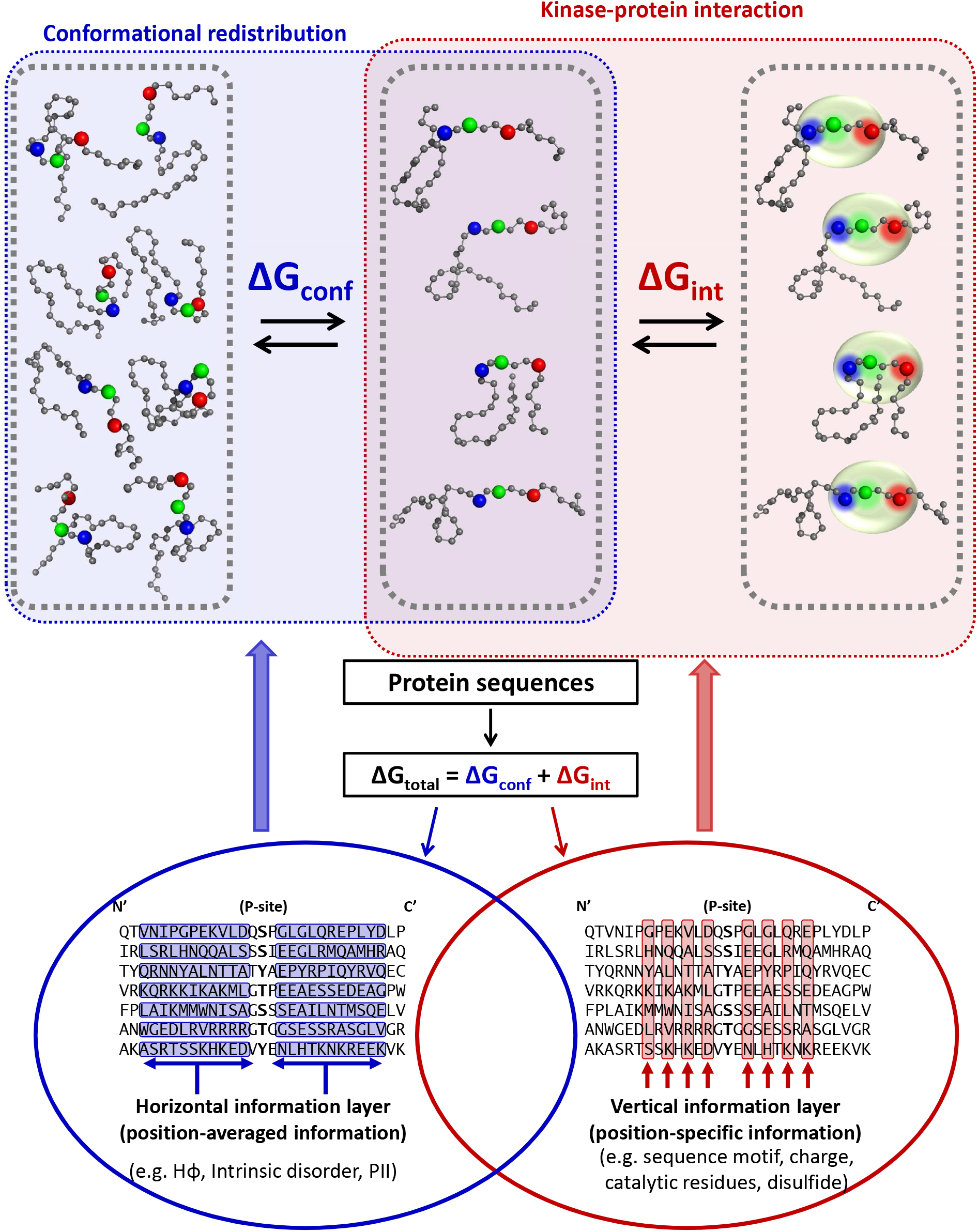
Horizontal and vertical protein sequence information reflected in the conformational and binding equilibria of kinase-substrate interaction. Cartoon of coupled equilibria (upper half) demonstrates a decrease of diversity in the substrate’s conformational ensemble mediated by horizontal information (blue box) necessary to position functional residues, mediated by vertical information (red box). Horizontal and vertical information are simultaneously encoded (lower half) in an amino acid sequence alignment. Black letters represent aligned sequences, with blue rows representing neighboring groups of amino acids exhibiting emergent biophysical properties, and red columns representing conserved amino acids typically used for alignment and binding site identification. The central hypothesis of this work is that biological phosphorylation, and effective phosphorylation site prediction, critically depends on both types of information.

In Expression (1), E is the kinase, S is the unphosphorylated substrate, and the subscripts BI and BC denote binding-incompetent and binding-competent conformations of the substrate. These equilibria are schematically depicted in Figure 1. Importantly, the binding-competent and binding-incompetent thermodynamic states are agnostic as to the degree of structure present, only that a free energy barrier exists between the sub-ensemble that can bind and be phosphorylated and the sub-ensemble that cannot.

Expression (1) defines two free energy contributions to protein phosphorylation: one from the organization of the intrinsically disordered substrate ensemble (Kconf) and one from binding of the organized substrate to the kinase active site (Kint). We hypothesize that these two contributions can be usefully separated and accessed in terms of quantifiable bioinformatics information (Fig. 1, red and blue circles).

In this scenario, both the substrate conformational ensemble and conserved recognition motif would encode the kinase specificity information, but the presence of two coupled equilibria might suggest two separate sources for this information. We define the ensemble based information as “horizontal”, meaning regionally distributed across a sequence fragment (Fig. 1, blue circle), while the conserved motif is “vertical”, meaning that the residue positions are largely independent (Figure 1, red circle). Importantly, the nature of these two types of information would suggest that horizontal information can be conserved even in the absence of significant vertical conservation.

### 2.2. Horizontal sequence information encodes conserved conformational dynamics

Our approach to accessing the residue–specific contributions to Kconf encoded in the horizontal information is predicated on previous results from our group showing that proteins can be represented as sequences of thermodynamic environments (21–23) that capture the experimental conformational fluctuations (24) in both ordered (25) and disordered (26) ensembles. We also showed that the propensities of amino acids in these thermodynamic environments provide sufficient information to match unknown sequences to their environmental profiles (23), and that these profiles are conserved (25, 27, 28) (see Supplemental Figure S10). The importance of these earlier findings is that they directly demonstrate that hidden information about the stability of a chain (reported at each position) is nonetheless embedded within the sequence, and can be accessed by comparing this horizontal information for diverse sequences, as schematically depicted in Figure 1 (Left).

Indeed, conservation of horizontal information may be even stronger than sequence conservation for some biological contexts, further motivating the combination of horizontal and vertical information. For example (Fig. 2A), conservation of the position-specific stability (31) among the members of the intrinsically disordered N-terminal domain of the glucocorticoid receptor family is high, while the amino acid conservation within the same domain is low (58). That such behavior seems to be a general feature of protein families (Fig. 2B) suggests that horizontal information is conserved to some degree even in the absence of amino acid conservation.

**Figure 2.**
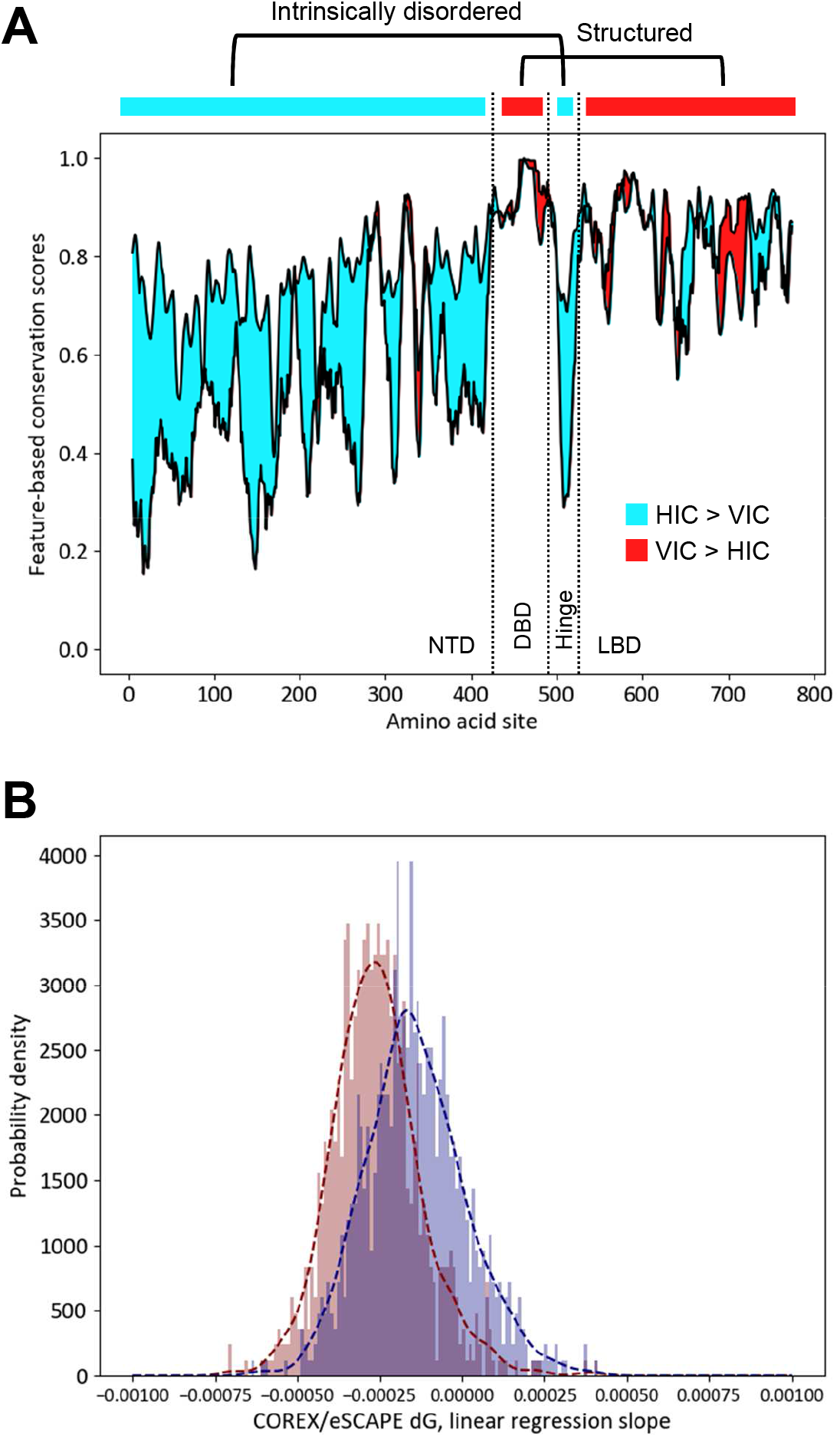
Horizontal information is more strongly conserved than vertical information in intrinsically disordered regions of protein families. A. Difference between degrees of conservation of sequence and native state free energy (∆G, (31)) calculated for human glucocorticoid receptor (GR) and its orthologs (58), Methods. Cyan denotes regions where free energy conservation (HIC, horizontal information conservation) is stronger than sequence conservation (VIC, vertical information conservation), and red denotes the opposite. In human GR, DNA binding region (DBD) and LBD region are structured, while N-terminal domain (NTD) and hinge region are intrinsically disordered. Preponderance of cyan area demonstrates that horizontal information can be conserved when vertical information is not. B. Coefficient of correlations between free energy and conservation score is calculated for ortholog alignments of 835 different transcription factors (58). Distribution of slope coefficients over many families show that sequence conservation (red) is more strongly correlated with calculated free energy, a property seen in Figure 2A for a single family.

### 2.3. Vertical sequence information from eukaryotic phosphorylation sites is distinguished primarily by the presence of +1 Pro

The classic approach to identifying phosphorylatable substrates has been to use independent position-conserved information (i.e. vertical information). To characterize the vertical information component, we investigated amino acid sequence fragments of 29 residues centered on known Ser, Thr, and Tyrosine (Tyr) eukaryotic phosphorylation sites (Fig. 3A). Immediately apparent from statistics of the human phosphoproteome is the abundance of Pro residues directly C-terminal to the annotated Ser or Thr phosphorylation site (Fig. 3A-B). Using the presence or absence of +1 Pro to separate phosphorylated and non-phosphorylated sequences into four subclasses reveals substantial differences in amino acid conservation patterns.

**Figure 3.**
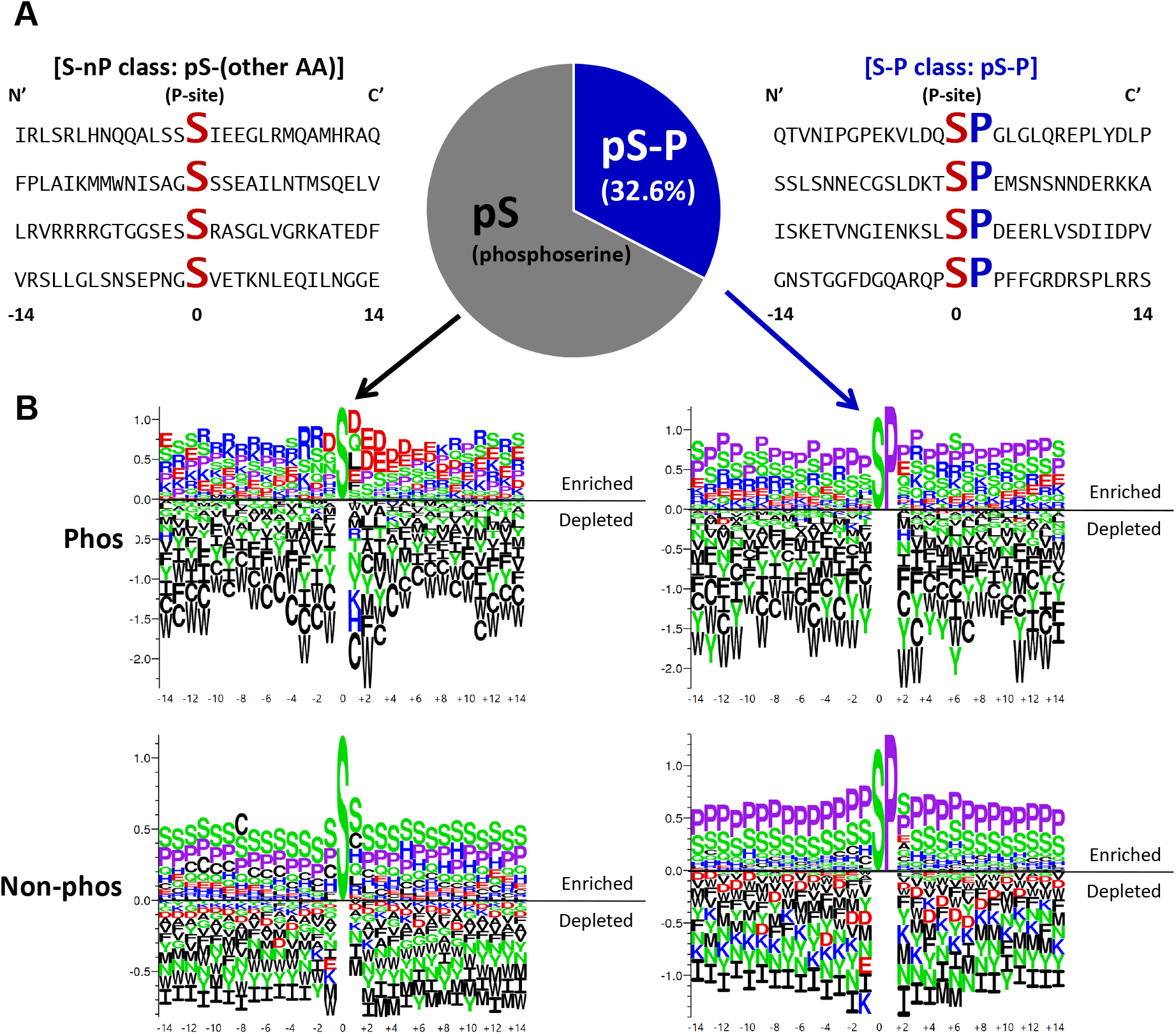
Proline residue at the +1 site (+1 Pro) of serine phosphorylation sites (pS) defines a subclass of site (pS-P) dependent on horizontal information. A. Example 29-mer sequence neighborhoods centered on the phosphorylated Ser residue. Conserved Ser (S) and +1 Pro residues (P) are enlarged and bold. Frequencies of +1 Pro phosphorylation sites (pS-P) make up one-third of all known human phosphorylated Ser. B. Amino acid frequencies around pS-P and pS-nP demonstrate that pS-P sites have little distinguishing sequence features as compared to S-P sites. Top logos show enrichment/depletion patterns of amino acids around phosphorylated Ser sites. Bottom logos show patterns around non-phosphorylated Ser sites. Left logos show patterns where the Ser is immediately followed by amino acids other than Pro. Right logos show patterns where the Ser is immediately followed by Pro (i.e. +1 Pro). Vertical scale indicates information content in bits.

Focusing on Ser sites as examples, all subclasses are generally depleted in hydrophobic and aliphatic residues (Fig. 3B). All subclasses except for phosphorylated non-(+1 Pro) sites are enriched with Ser and Pro, implying enrichment of intrinsic disorder (Supplementary Figure S1). In contrast, phosphorylated non-(+1 Pro) sites exhibit enrichment of positively charged amino acids Arginine (Arg) and Lysine (Lys) at positions N-terminal to the phosphorylation site, and enrichment of negatively charged amino acids Aspartate (Asp) and Glutamate (Glu) at positions C-terminal to the site (Fig. 3B, top left), distinguishing the sequence neighborhoods of phosphorylated and non-phosphorylated sites. Sites with +1 Pro are more difficult to distinguish based on sequence conservation alone, although the phosphorylated sites appear to tolerate a certain amount of Glu (Fig. 3B, top right) while the non-phosphorylated sites are depleted in all negatively charged side chains (Figure 3B, bottom right).

Surprisingly, when the presence of the +1 Pro is ignored, the sequence neighborhoods of Ser phosphorylated +1 Pro sites (Fig. 3B, top right) are similar to those of non-phosphorylated non-(+1 Pro) sites (Figure 3B, bottom left), with both subclasses enriched in Pro, Ser, and Glu. This indicates that there is little conserved sequence information to locally distinguish a phosphorylated site from a non-phosphorylated one. Indeed, inspection of the logos suggests that Ser phosphorylation sites, for example, are especially depleted in aromatic amino acids (Figure 3B, top) relative to non-phosphorylated sites (Figure 3B, bottom). Simple positional conservation would report the absence of aromatics at all sites, but experimental results demonstrate that even single aromatic substitutions in an otherwise identical background could have large effects on denatured state properties (29).

Many classic examples of vertical information used in phosphorylation site prediction have been previously reported (10, 13, 14, 16), but we focus here on the special case of Ser and Thr sites with +1 Pro (noting Tyr shows no +1 sites), and demonstrate that this sequence motif, although not diagnostic by itself, is particularly useful in site prediction. Testing several residue types and locations in the neighborhood of known phosphorylation sites, the presence of +1 Pro is the single most informative position in differentiating subgroups from the complete dataset (Supplementary Figure S2B). Thus, we can partition sites into five subclasses based on the presence or absence of the +1 Pro at Ser and Thr phosphorylatable sites: Ser +1 Pro (S-P), Thr +1 Pro (T-P), Ser non-(+1 Pro) (S-nP), Thr non-(+1 Pro) (T-nP), Tyr (Y) (Supplementary Figures S2C-D). This grouping is supported by the observation that position-specific weight matrices constructed from these subclasses are more similar between Ser +1 Pro and Thr +1 Pro than between either Ser non-(+1 Pro) and Ser +1 Pro or Thr non-(+1 Pro) and Thr +1 Pro (Supplementary Figure S2C). For consistency with previously published work from other researchers, we also considered the simpler three subclass grouping based only on the identity of the phosphorylatable residue (Supplementary Figure S2D).

### 2.4. Phosphorylatable sites in disordered substrates with +1 Pro are poised to respond to the phosphorylation event

Accepted sequence heuristics exist that map expected conformational states of folded and disordered protein sequences to their PII propensity (20, 30), conformational stability (23, 26, 31–34, 56), or polarity and charge properties (35, 36). Given the demonstrated importance of +1 Pro in phosphorylation site subclass identification, we sought to understand the influence of these effects on the conformational manifolds of the subclasses. In particular, we predicted end-to-end ensemble distances (30) (Fig. 4A) and mapped annotated phosphorylation sites to the charge-charge plots of the denatured state (Fig. 4C) introduced by Das, et al. (35) to understand the expected conformational properties before and after the phosphorylation event (see Methods).

**Figure 4.**
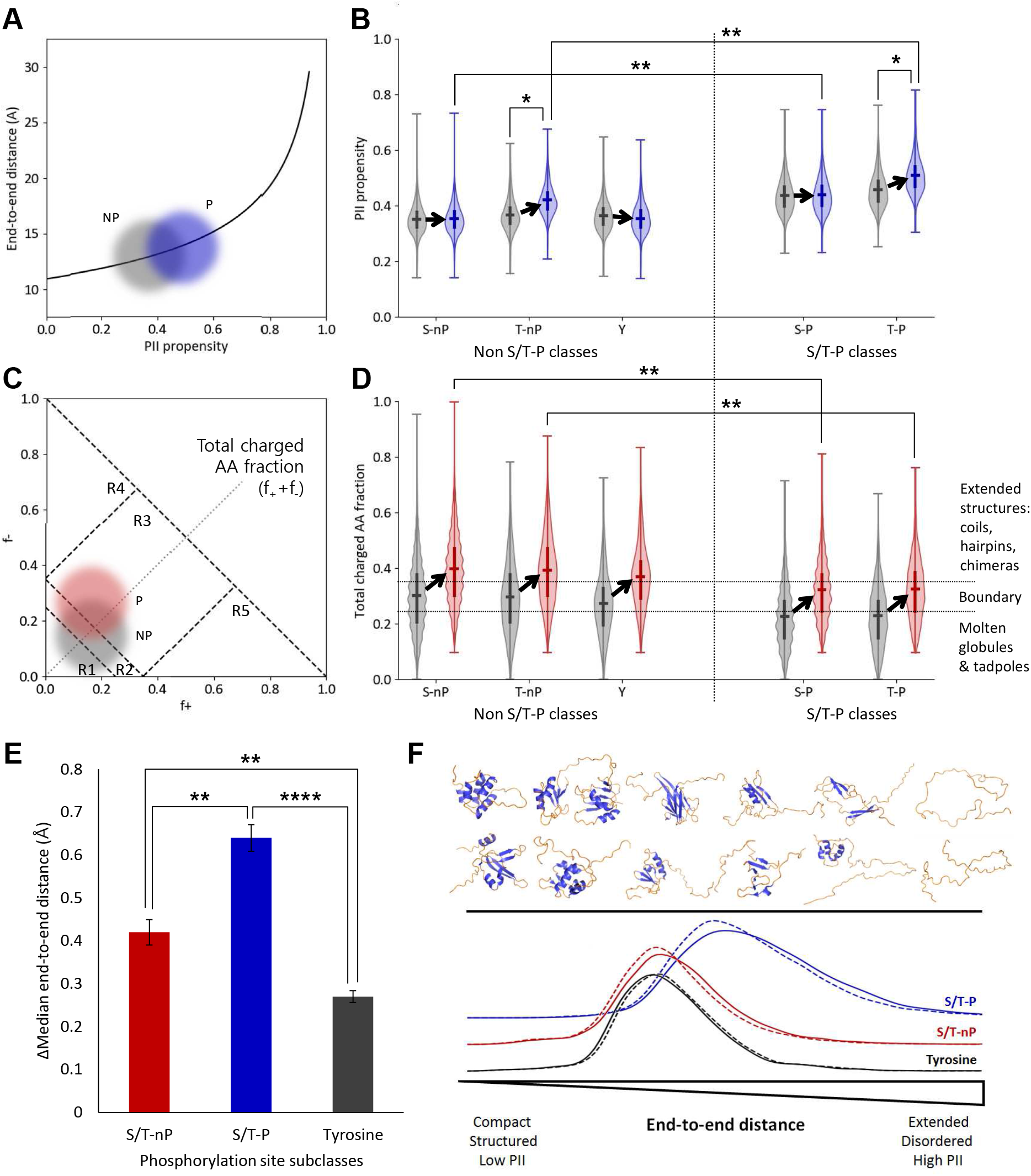
Phosphorylation sites containing +1 Pro are energetically poised to respond to phosphorylation by extension, mediated by charge and PII propensity. A. Conceptual plot illustrating expected end-to-end distance increase (30) due to phosphorylation of an ensemble distribution of 29-mer sequence fragments. Gray cloud represents non-phosphorylated sequences (NP) and blue cloud represents singly-phosphorylated sequences (P). B. Violin plots of ensemble distributions of sequence PII propensities (20) before (gray) and after (blue) phosphorylation. The +1 Pro classes in particular (the two right-most pairs of distributions) exist in an extension range nearest the exponential increase in panel A. C. Conceptual plot illustrating expected charge change due to single phosphorylation (P) of a distribution of 29-mers. The numbered regions R1-R5 represent conformational regimes as described in Das, et al. (35). Note that the dashed diagonal line corresponds to the y-axis in panel D, following. D. Violin plots of ensemble distributions of sequence charge properties before (gray) and after (red) phosphorylation. Dotted horizontal lines represent conformational regimes as described in Das, et al. (35). Sites with +1 Pro (the two right-most pairs of distributions) specifically exhibit a less unstructured conformational manifold prior to a phosphorylation event, thus the Pro effectively buffers a conformational transition with an increased PII propensity. E. +1 Pro sites undergo the largest expected extension upon phosphorylation due to contributions from both extension (PII structure) and charge repulsion (see Supplementary Figures S3-6). F. Schematic summarizing changes in the conformational ensemble upon phosphorylation. The top-half represents an idealized conformational spectrum ranging from mostly folded (left side) to mostly disordered (right side). Conformational change is measured by end-to-end distance (bottom), mediated by PII propensity and charge interactions. Along this spectrum, tyrosine phosphorylation (gray arrow) exhibits the smallest change, non +1 Pro site phosphorylation (red arrow) exhibits a moderate change, and +1 Pro site phosphorylation exhibits the largest change (blue arrow).

Distributions of the sequence properties for each of the five subclasses suggest not only that the computed dimensions of respective conformational ensembles are poised in statistically different regions, but that the different subclasses respond differently to a phosphorylation event, with the +1 Pro sites responding more similarly than the non-(+1 Pro) sites (Figs. 4B and 4D). In detail, the generally expected response to adding the extra negative charge of a phosphoryl group is to make the sequence neighborhood less structured (Fig. 4C) (35), due both to the energetic unfavorability of burying a charged sequence in a structured hydrophobic core and to the energetic unfavorability of other like charges in the sequence neighborhood. Although the distribution is broad, this expected behavior is seen for the Ser and Thr non-(+1 Pro) subclasses (Figs. 4B and 4D, first two pairs of distributions).

The +1 Pro subclasses exhibit a different response, becoming less unstructured than expected (Fig. 4B and 4D). However, the higher Pro content of these subclasses, and the high predicted sequence disorder content, do not support a folding event for the +1 Pro subclass. In fact, this subclass exhibits the highest increase in end-to-end distance (Fig. 4E), despite the fact that this increase starts from a more unstructured conformational ensemble. We hypothesize that modulating the population of PII structure through phosphorylation expands the ensemble (20), and that sequences already high in PII are more sensitive to the effects, as Fig. 4A would suggest. The demonstrated ability of Ser/Thr phosphorylation to selectively increase end-to-end distance in high PII sequences (20, 37) supports this hypothesis.

### 2.5. An improved phosphorylation site predictor resulting from consideration of conformational dynamics

To explore the practical manifestations of our findings, the horizontal and vertical information were incorporated into a novel prediction method called PHOSforUS (see Methods). Evaluation of the individual predictors with cross-validation demonstrates that all five subclass-specific predictors have good predictive power for identifying annotated phosphorylation sites (Table 1). The horizontal and vertical information-specific predictors have similar performance, with the horizontal combination, including the position-specific COREX information (23, 26, 31–34) contained in eScape, showing the best performance (Figure 5B, blue curve). Although combining both horizontal and vertical information results in improved prediction accuracy relative to either alone (Figure 5B, black curve), horizontal information consistently is more effective (as measured by AUROC) across subclasses than vertical information alone (Figure 5C, Supplementary Figures S7-9 and Supplementary Table 9). This result indicates that conformational equilibrium is the most important contributor to the phosphorylation event.

**Table 1.**
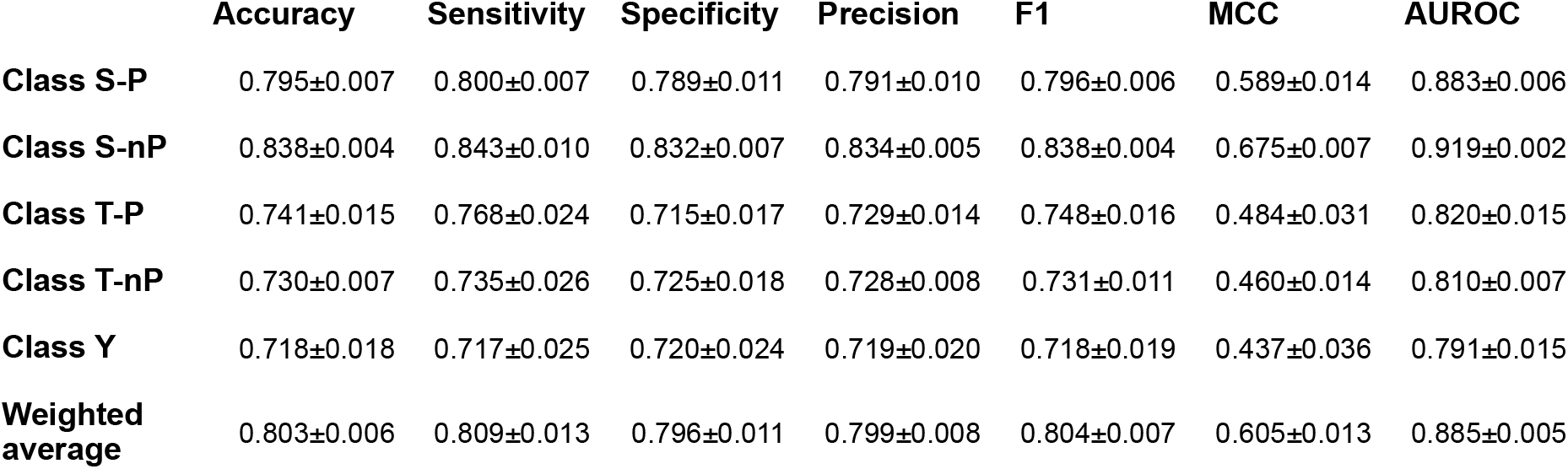
Subclass training performance of *PHOSforUS* predictor.

**Figure 5.**
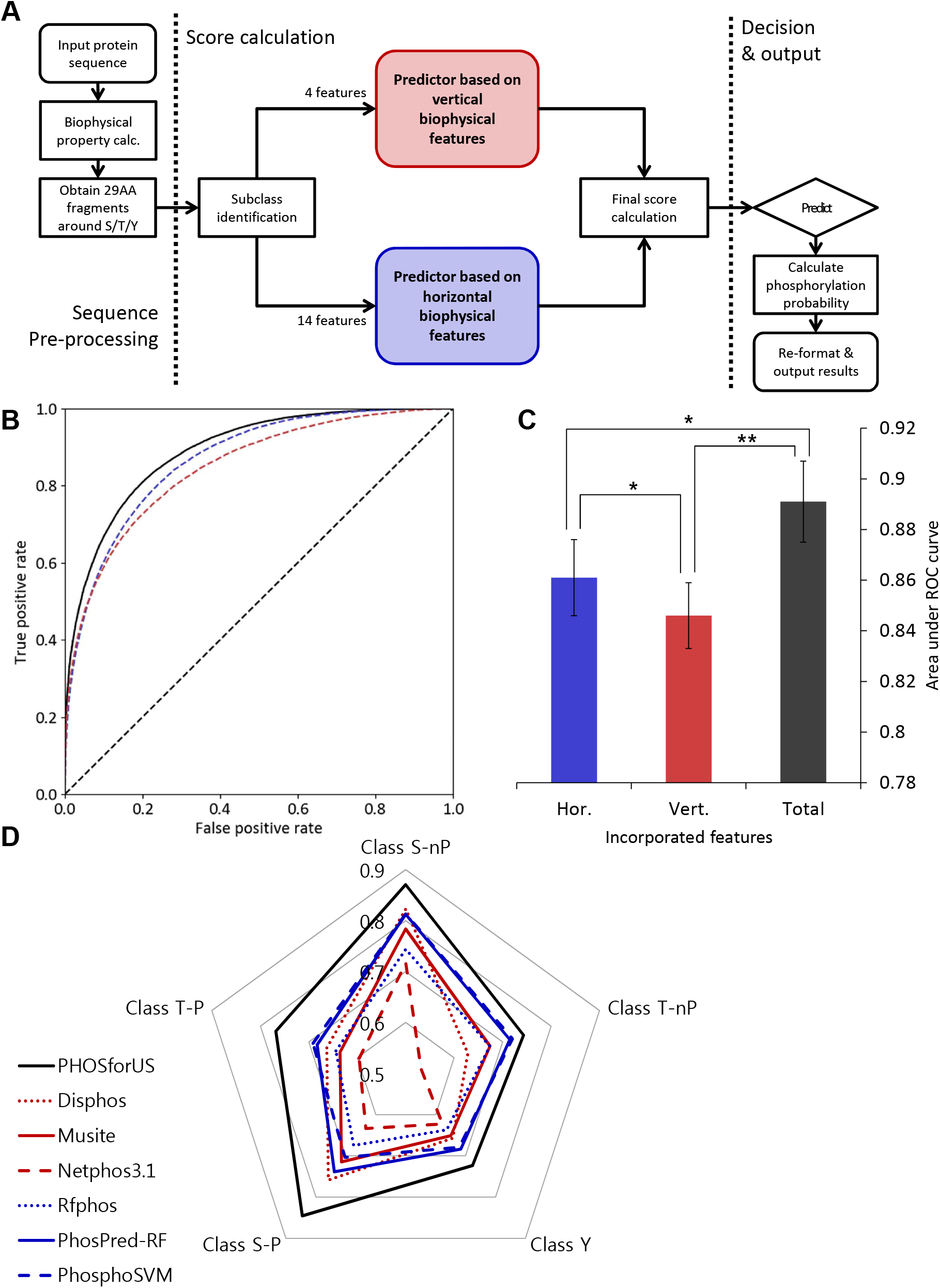
Architecture, training performance, and comparative effectiveness of PHOSforUS predictor. A. Simplified workflow of PHOSforUS predictor algorithm. Biophysical properties of an arbitrary protein sequence are split into 29-mer fragments centered on Ser/Thr/Tyr residues. Five (or three) subclass-specific predictors are invoked, independently based on vertical (red) or horizontal (blue) information. Intermediate output is combined with gradient boost, and combination scores over a preset threshold are predicted as phosphorylated. B. Receiver–operating characteristics (ROC) of PHOSforUS constituent predictors. Area under the ROC curve (AUROC) is indicated as a separate bar graph (C). Performance of all subclasses of phosphorylation site are combined into a single curve. The combined predictor (Total, black) outperforms separate predictors based on vertical (Vert, red) or horizontal (Hor, blue) information. Notably, horizontal information significantly outperforms vertical information (C), demonstrating the importance of horizontal information. D. Comparative effectiveness of protein phosphorylation site prediction by PHOSforUS. For five subclasses of phosphorylation site, PHOSforUS AUROC values meet or exceed those obtained on the identical data with six existing prediction tools.

Although several dozen prediction methods exist (16), six available tools were compared with PHOSforUS to assess the algorithm’s real-world performance (10, 11, 13–16). These methods were chosen because they were freely accessible and could handle the large datasets used for testing (see Methods). Based on ROC curves, the seven methods broadly segregated into two groups, with the most effective group containing methods that either explicitly, or implicitly, incorporated disorder prediction information (Table 2). For all five site classes, PHOSforUS exhibited the highest AUROC values (Fig. 5D and Table 2). Because we cannot exclude the possibility that phosphorylation sites in the testing set were not already contained in the training sets for the other methods, it is likely that the performance improvement of PHOSforUS reported here is a conservative estimate.

**Table 2.** Comparative analysis (AUROC) of PHOSforUS against currently utilized predictors.

| **Subclass** | ***PHOSforUS*** | ***Disphos*** | ***Musite*** | ***Netphos3.1*** | ***Rfphos*** | ***PhosPred-RF*** | ***PhosphoSVM*** |
| --- | --- | --- | --- | --- | --- | --- | --- |
| **Class S-nP** | 0.871±0.028 | 0.823±0.028 | 0.783±0.037 | 0.717±0.044 | 0.744±0.026 | 0.813±0.034 | 0.814±0.025 |
| **Class T-nP** | 0.743±0.018 | 0.628±0.018 | 0.674±0.019 | 0.531±0.042 | 0.674±0.036 | 0.715±0.029 | 0.720±0.024 |
| **Class Y** | 0.724±0.024 | 0.657±0.024 | 0.651±0.039 | 0.622±0.022 | 0.637±0.067 | 0.684±0.022 | 0.678±0.021 |
| **Class S-P** | 0.845±0.046 | 0.758±0.046 | 0.715±0.030 | 0.633±0.053 | 0.674±0.032 | 0.738±0.027 | 0.703±0.054 |
| **Class T-P** | 0.768±0.023 | 0.663±0.023 | 0.635±0.011 | 0.596±0.033 | 0.644±0.037 | 0.683±0.023 | 0.692±0.030 |
| **Weighted average** | 0.836±0.021 | 0.767±0.030 | 0.738±0.032 | 0.665±0.044 | 0.707±0.032 | 0.769±0.030 | 0.762±0.018 |

The implications of this result are three-fold. First, the prediction effectiveness of PHOSforUS is evidence that the ensemble conformational transition is important for kinase recognition. Second, because PHOSforUS is trained on sequence fragments, information about the conformational dynamics (i.e. fluctuations) of the phosphorylation site is contained in the sequence neighborhood of the site. Third, because the +1 Pro can meaningfully segregate human phosphorylation sites, it is possible that phosphorylation site sequence logos (Supplementary Figures 1 and 2) can in some cases also reflect evolutionary conservation of horizontal information.

## 3. Discussion

Here we tested whether embedded horizontal information that captures the thermodynamics of a disordered chain plays a role in determining the ability of a substrate to be phosphorylated. Yet, phosphorylation can be considered as a specific case of a more general problem: how to identify the structural determinants of any biochemical reaction targeted to an intrinsically disordered site. Our proposed solution is to reformulate the problem of site prediction, whether structured or disordered, by couching it within a thermodynamic framework. In the case of a target site within a structured protein, such a framework is simplified by the existence of a unique native structure. In the case of intrinsic disorder, we employ a “thermodynamic proxy” due to the absence of accurate conformational ensemble modeling technology (though such technology is rapidly developing (38–41)).

In this work, we schematically represented the phosphorylation reaction and phosphorylation prediction in terms of two distinct processes with their corresponding free energies (Fig. 1). The “vertical” information is reflective of the classic static structural view of proteins and substrates, whereby the conserved sequence elements provide the scaffold for tight binding. In effect, the degree of conservation serves as a proxy for the energy of the interaction, a result that is consistent with the reported similarity in statistical vs. experimental energy changes observed within folded proteins (60).

Unique to the approach described here, however, is the incorporation of “horizontal” information that specifically encodes the conformational free energy differences embedded along a sequence. Importantly, both types of information could be encoded by amino acid sequence and should be conserved in a substrate multiple sequence alignment (Fig. 1, circles), with a key difference being that the horizontal information is more diffuse and thus would be expected to be less conserved at individual positions using traditional alignment tools (42). This could be an indication of an evolutionary strategy that permits rapid testing of functional amino acid substitutions within a conserved disordered region. Support for the relevance of horizontal information comes from the direct comparison of sub-predictor statistics, such as area under ROC curve and accuracy, which reveals that horizontal features perform better than vertical features in every phosphorylation subclass (Supplementary Tables 4-8).

The presence or absence of the +1 Pro is a key feature for subclass identification and for the effectiveness of PHOSforUS predictions. What is the biological function of a phosphorylated side chain followed by a Pro, and why is the +1 Pro motif common in eukaryotes and not prokaryotes? Although speculative, our results suggest the answer lies in the work that is done in the form of conformational extension upon phosphorylation. To appreciate this point, it must be remembered that there are at least two documented mechanisms for extension in a disordered ensemble: changes in charge mixing (Fig. 4C) (35, 36, 38) and changes in intrinsic polyproline II propensity (Figure 4A) (20, 30, 36, 37). Sequence logos (Supplementary Figure S1) demonstrate that the second mechanism is likely to be associated with +1 Pro subclass.

To assess the relative extension for these phosphorylated sequence fragments, we used the method of Tomasso and colleagues (30), which takes both charge and PII propensity into account. Phosphorylation subclasses with +1 Pro show a significant extension expected post-phosphorylation of more than 0.6 Å for a 29-mer (Figure 4E). Notably, this extension is mediated by both charge and PII propensity (Supplementary Figures S3-6). Thus, +1 Pro sites may encode a conformational switch between disorder and PII extended structure. Detailed investigation of individual proteins will be required to ascertain the biological functions of such a switch, but we speculate that some cases are related to the efficiency of the signal transmitted by the phosphoryl label, while others could be related to the emerging phenomenon of liquid droplet stress granule formation in eukaryotes (43).

## 4. Conclusion

We have shown that horizontally conserved information regarding the structure and energy of the conformational ensemble of a protein sequence plays a major role in determining which disordered sequences will be phosphorylated and how these ensembles will be affected by phosphorylation. Importantly, we note the model presented here is not a rigorous statistical thermodynamic method that explicitly accounts for specific interactions and contributions of individual amino acids. Instead, we asked whether the hidden conformational free energy information previously demonstrated to be embedded within all protein sequences, is sufficient to provide predictive information when sequence conservation is too low to render meaningful comparisons. Our ability to take as input a single amino acid sequence and predict the likelihood of phosphorylation at Ser, Thr, and Tyr residues (Figure 5), demonstrates the validity of our approach, and supports our assertion that conformational dynamics (or fluctuations) can affect (and in the case of phosphorylation, even dominate) the specificity of a biological process. Thus in some respects our development of a state-of-the-art prediction algorithm can be viewed as a “byproduct” (albeit highly desirable) of the more important biological finding, which demonstrates the critical role played by conformational dynamics in determining the functional regulatory changes in intrinsically disordered proteins.

## 5. Materials and Methods

### 5.1. Reference dataset and data processing

Canonical human protein sequences were obtained from SWISS-PROT (2018 December Release) (8), a manually curated subset of the UniProt database. Phosphorylation annotations were obtained from SWISS-PROT and PhosphoSitePlus (2018 December Release) (9). True positive sets were assembled from SWISS-PROT annotations and low-throughput (LTP) subset of PhosphoSitePlus. Sequence fragments of 29 amino acids (14 residues N-terminal and C-terminal relative to a central phosphorylation site) were extracted from these sets and subsequently divided into five subsets (S-P, S-NP, T-P, T-NP, Y) based on the identity of the center residue and the presence of Pro as its C-terminal neighbor. For example, S-P denotes Ser as the phosphorylatable central residue with presence of the +1 Pro, while S-NP denotes any of the remaining 19 residues at the +1 position. To reduce information redundancy, a 50% maximum pairwise sequence similarity filter was applied to these subsets. True negative subsets were assembled in a similar way and sequences that shared more than 50% similarity to any phosphorylated sequence were removed to filter out false positives. Resulting statistics of these sets are shown in Supplementary Table S1.

For the comparative analysis, we constructed another positive set which contains none of the sequences already contained in the training set, and presumably minimal number of sequences in the training sets of existing phosphorylation predictors. From PhosphoSitePlus high-throughput (HTP) subset, we removed sequences that show 50% similarity to any of sequences within SWISS-PROT, Phospho.ELM (7) and PhosphoSitePlus LTP datasets.

From resulting positive set (statistics shown in Supplementary Table S1) and true negative set, we randomly sampled 5 testing sets with 100 positive sites and 100 negative sites to test predictor performances.

### 5.2. Visualizing conservation of vertical and horizontal information

Orthologs of human proteins with DNA-binding transcription factor activity (GO: 0003700) were obtained from OMA database (58). We selected ortholog groups with the number of members between 10 < n < 250, and downloaded multiple sequence alignments as archived in the database. A full list of the 835 ortholog groups we utilized is found in Supplementary Data File 2.

Normalized local sequence conservation scores were calculated using the following procedure. The multiple sequence alignment (alignment size = n) was divided into small overlapping windows (window size = 5, step = 1). For each window, pairwise local alignment scores using BLOSUM62 matrix (59) were calculated between a reference sequence (*Seq*_*i*_) and each of all other sequences within same ortholog group (*Seq*_*j*_). This process was iterated using each of the sequences in the alignment as a reference sequence. Within each iteration, each pairwise score was divided by a maximum score attainable, which was defined as the case when a sequence which is identical to the reference was applied for pairwise comparison. Calculated pairwise scores were averaged to obtain a normalized local sequence conservation score (Equation 2).

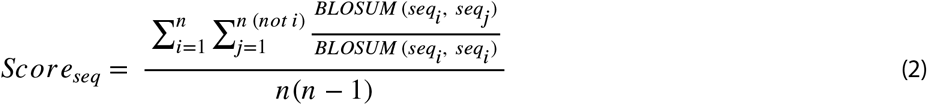

Native state free energy for each protein sequences was calculated using the eSCAPE algorithm (31, https://best.bio.jhu.edu/eScape). For the same window we used for calculation of local sequence conservation score, we calculated local average and standard deviation of free energy values. Horizontal conservation score was computed using the following Equation 3:

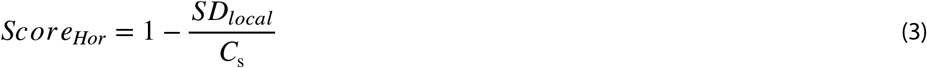

In this case, scaling coefficient (*C*_s_ = 3.3 (kcal/mol)) was calculated from 10 different ortholog groups exhibiting high sequence conservation and structural stability (for example, actin (ACTB) and rhodopsin (RHO) families). Resulting conservation scores are plotted in Figure S10A (glucocorticoid receptor / GCR), Supplementary Figure S10B (actin) & S10C (rhodopsin) respectively.

To observe its correlation with free energy, sequence conservation scores and horizontal conservation scores were normalized again with μ = 0 and SD = 1 (i.e. a Z-score). Linear correlations between average free energy and both conservation scores were calculated subsequently as Figure S10D. Binned distributions for slopes and correlation coefficients (for 835 correlations, one for each ortholog group) could be found in Figure S10E and S10F respectively.

### 5.3. Combining horizontal and vertical information to build a phosphorylation site predictor

Selected horizontal information was computed over 29 residue window (Supplementary Information) using properties contained within the AAindex database (44). Properties that were not classified as horizontal were considered vertical information. A naïve Bayes predictor (45, 57) trained on each individual property was used to assess predictive accuracy for each phosphorylation subclass (Supplementary Tables S4-S8), and the individual properties with highest information content were incorporated into the PHOSforUS prediction algorithm (Supplementary Tables S2 and S3). Horizontal properties included amino acid partition energies (46, 47), alpha helix frequencies (48), extended conformation (49, 50) and polyproline II helix propensities (20), hinting at cooperative and non-cooperative structure tendencies. Vertical properties included amino acid isoelectric point (51), molecular weight (52), volume (53), and side chain average exposed surface area (54), all being characteristics independent of neighboring amino acids. Orthogonal information was incorporated from predicted thermodynamic properties (23, 26, 31–34) using the eScape software (31), this information was used to train a separate naïve Bayes predictor (Supplementary Information).

The PHOSforUS algorithm consisted of three stages: sequence pre-processing, score calculation, and decision output (Figure 5A). The first stage identifies the Ser, Thr, and Tyr residues as possible phosphorylation sites and computes the horizontal and vertical properties mentioned above for each site’s sequence neighborhood. The second stage routes each site to the appropriate subclass predictor and parameter set. Prediction scores from each individual horizontal, vertical, and thermodynamic property are combined using a Gradient Boost (55, 57) predictor (see details of predictor architecture, Supplementary Information), resulting in a single value for the potential site. The third stage compares this single value to a pre-determined threshold to predict the probability that the site is phosphorylated or non-phosphorylated. Thus, a confidence is attached to the binary phosphorylation prediction, making the prediction more interpretable to the researcher.

### 5.4. PHOSforUS source code, software, and databases

The PHOSforUS software package and associated databases are freely available at https://github.com/bxlab/PHOSforUS.

## Supporting information

Supplementary Data File 2: List of the ortholog groups

## 6. Acknowledgements

Funding from NIH (R01-GM063747, U41 HG006620), NSF (MCB-1330211), and Johns Hopkins University is gratefully acknowledged.

