## Supplementary Text, Figures, and Tables for "Hidden dynamic signatures drive substrate selectivity in the disordered phosphoproteome"

* Vincent J. Hilser or James Taylor

**This PDF file includes:**

Supplementary text

Figures S1 to S10

Tables S1 to S10

SI References

### Supplementary Information Text

*Details of feature selection and predictor architecture*

Features were selected from a pool of 546 amino acid scales based on information content. These scales were obtained from the *AAindex* database (1) with the addition of the *DisProt* (2) and experimental poly-Proline II (PII) (3) propensity scales. To assess information content of a scale, analysis datasets were built with randomly selected 1000 true positives and 1000 true negatives, both coming from the same phosphorylation subset of 29-residue fragments. For every fragment, a weighted average of values from the scale with window size nine was calculated for the region of 21 residues centered on the Ser/Thr/Tyr, and the information value of the scale was estimated using a naïve Bayes classifier (4) with ten-fold cross validation. This procedure was iterated ten times with randomly selected analysis datasets and tested for each of the five phosphorylation subclasses individually. For each subclass, an amino acid scale that exhibited greater than 0.6 prediction accuracy was retained, otherwise the scale was rejected. In this way, 114 scales were retained from the original pool. This number was further reduced to 35 by only retaining one member of scale pairs exhibiting an absolute Pearson correlation coefficient greater than 0.8. Finally, manual curation to remove redundant scales based on AAindex descriptors resulted in ten features used in the predictor (Supplementary Table S2).

Additional features for the predictor were obtained from our unique sequence-based energy prediction tool, *eScape* (5, 6). This tool, originally parameterized using tripeptide-based protein ensemble energetics (5), predicts stability, enthalpy, and entropy of both native and denatured states by using 28 feature indices. From these 28 features, we empirically selected four native state features and four denatured state features whose values and differences seemed to be effective in phosphorylation site prediction. The eight features and differences are listed in Supplementary Table 3. Thus, a total of 18 features were used in the final predictor, and the whole list of features and parameters employed are shown in Supplementary Table 4.

The architecture of the *PHOSforUS* predictor is shown in Figure 5A. From an arbitrary input amino acid sequence, 18 biophysical features (Supplementary Tables 2 and 3) are calculated for each 21-residue fragment of the sequence centered on Ser, Thr, or Tyr residues. Thus, for each Ser/Thr/Tyr residue, the +1 Pro subclass is assigned and a total of 378 feature values are calculated (21 residue positions x 18 biophysical features).

Information values for each potential phosphorylation site were calculated from sub-predictors corresponding to each calculated feature value. Sub-predictors are Pro-subclass specific and are based on Gaussian naïve Bayes classifier (4, 7) (Equation S1). For each feature $F_i = \{f_{i1}, \ldots, f_{in}\}$, a final sub-predictor score $s_i$ is calculated as:

$$s_i = P\left(C_{phos} \middle| \{f_{i1}, \ldots, f_{in}\}\right) = \prod_{j=1}^{n} \frac{P\left(f_{ij} \middle| C_{phos}\right) P\left(C_{phos}\right)}{P\left(f_{ij}\right)} \quad . \tag{S1}$$

This intermediate output of sub-scores is passed to a downstream meta-predictor based on gradient boosting classifier (7, 8), which utilized those values to compute a final prediction score for each potentially phosphorylatable residue (Equation 3). For a set of sub-scores $S = \{s_1, \ldots, s_n\}$, where $n$ = 18 features, the score function $f(S)$ is fit with:

$$f(S) = f_0(S) - \sum_{j=1}^{m} \gamma_j \sum_{i=1}^{n} \nabla_{f_{j-1}} L\left(y_i, f_{j-1}(S_i)\right) \quad . \tag{S2}$$

In Equation (S2), the $\gamma_j$ term is equal to:

$$\gamma_j = arg \min_{\gamma} \sum_{i=1}^{n} L\left(y_i, f_{j-1}(S_i) - \gamma \nabla_{f_{j-1}} L\left(y_i, f_{j-1}(S_i)\right)\right) \quad . \tag{S3}$$

Phosphorylation likelihood was converted from the final score (Equation S2) and re-formatted for machine-readable output,

$$P\left(C_{phos}|input\ sequence\right) = \frac{P\left(f(S)|C_{phos}\right)P\left(C_{phos}\right)}{P\left(f(S)\right)} \quad . \tag{S4}$$

*Subclass prediction model evaluation*

10-fold cross-validation was performed to evaluate the sensitivity, specificity, and accuracy of the prediction models. As the true negative set is much larger than the true positive set, random sampling of the true negative set equalized the numbers of true and false positives during the evaluation. Cross-validation was iterated ten times with different true negative sets to minimize sampling error.

*Comparative analysis*

*NetPhos2.0* (9), *Musite* (10), *DisPhos* (11), *PhosphoSVM* (12), *RF-Phos* (13), and *PhosPred-RF* (14) were used to benchmark *PHOSforUS*. For the comparative analysis, we constructed another positive set which contains none of the sequences already contained in the training set, and presumably minimal number of sequences in the training sets of existing phosphorylation predictors. Details of how we prepared testing set are elaborated in the Main Text, Methods.

The following evaluation metrics were used: True Positive Rate (Equation S5), True Negative Rate (Equation S6), Positive Predictive Value (Equation S7), Accuracy (Equation S8), F1 Score (Equation S9), and Matthews Correlation Coefficient (Equation S10). In Equations (S5) – (S10), *TP* stands for true positive, *FN* for false negative, *FP* for false positive, *TN* for true negative.

$$\text{Sensitivity} = \frac{TP}{TP+FN} \tag{S5}$$

$$\text{Specificity} = \frac{TN}{TN+FP} \tag{S6}$$

$$\text{Precision} = \frac{TP}{TP+FP} \tag{S7}$$

$$\text{Accuracy} = \frac{TP+TN}{TP+FN+FP+TN} \tag{S8}$$

$$\text{F1 score} = \frac{2TP}{2TP+FP+FN} \tag{S9}$$

$$\text{MCC} = \frac{TP\times TN - FP\times FN}{\sqrt{(TP+FP)(TP+FN)(TN+FP)(TN+FN)}} \tag{S10}$$

*Visualizing conservation of vertical and horizontal information*

Orthologs of human proteins with DNA-binding transcription factor activity (GO: 0003700) were obtained from OMA database (17). We selected ortholog groups with the number of members between 10 < n < 250, and downloaded multiple sequence alignments as archived in the database. A full list of the 835 ortholog groups we utilized is found in Supplementary Data File 2.

Sequence conservation scores were calculated by using BLOSUM62 matrix (18). Single sequence was taken from an ortholog group as a reference and divided into small windows (window size = 5). For each window, pairwise local alignment scores were calculated between the reference sequence and each of all other sequences within same ortholog group, then all scores were divided by the maximum possible score $S_c$ (defined as the score calculated with identical sequence to the reference). This process was repeated for all other sequences within the ortholog group and averages over each window were taken as sequence conservation scores. Native state free energy for each protein sequences was calculated using the *eSCAPE* algorithm (5, https://best.bio.jhu.edu/eScape). For the same window we used for calculation of sequence conservation score, we calculated local average and standard deviation of free energy values. Horizontal conservation score was computed using the following Equation S11:

$$Score_{Hor} = 1 - \frac{SD_{local}}{S_c} \tag{S11}$$

In this case, scaling coefficient ($S_c =$ 3.3 (kcal/mol)) was calculated from 10 different ortholog groups exhibiting high sequence conservation and structural stability (for example, actin (ACTB) and rhodopsin (RHO) families). Resulting conservation scores are plotted in Supplementary Figure S10A (glucocorticoid receptor / GCR), Supplementary Figure S10B (actin) & S10C (rhodopsin), respectively.

To observe its correlation with free energy, sequence conservation scores and horizontal conservation scores were first normalized again with μ = 0 and SD = 1 (*i.e.* a Z-score). Linear correlations between average free energy and both conservation scores were calculated subsequently: slope values and Pearson correlation coefficients were collected for further statistical analysis. Collected slopes for 835 correlations, one for each ortholog group, are displayed as binned distributions in Supplementary Figure 10E.

**Supplementary Figures**

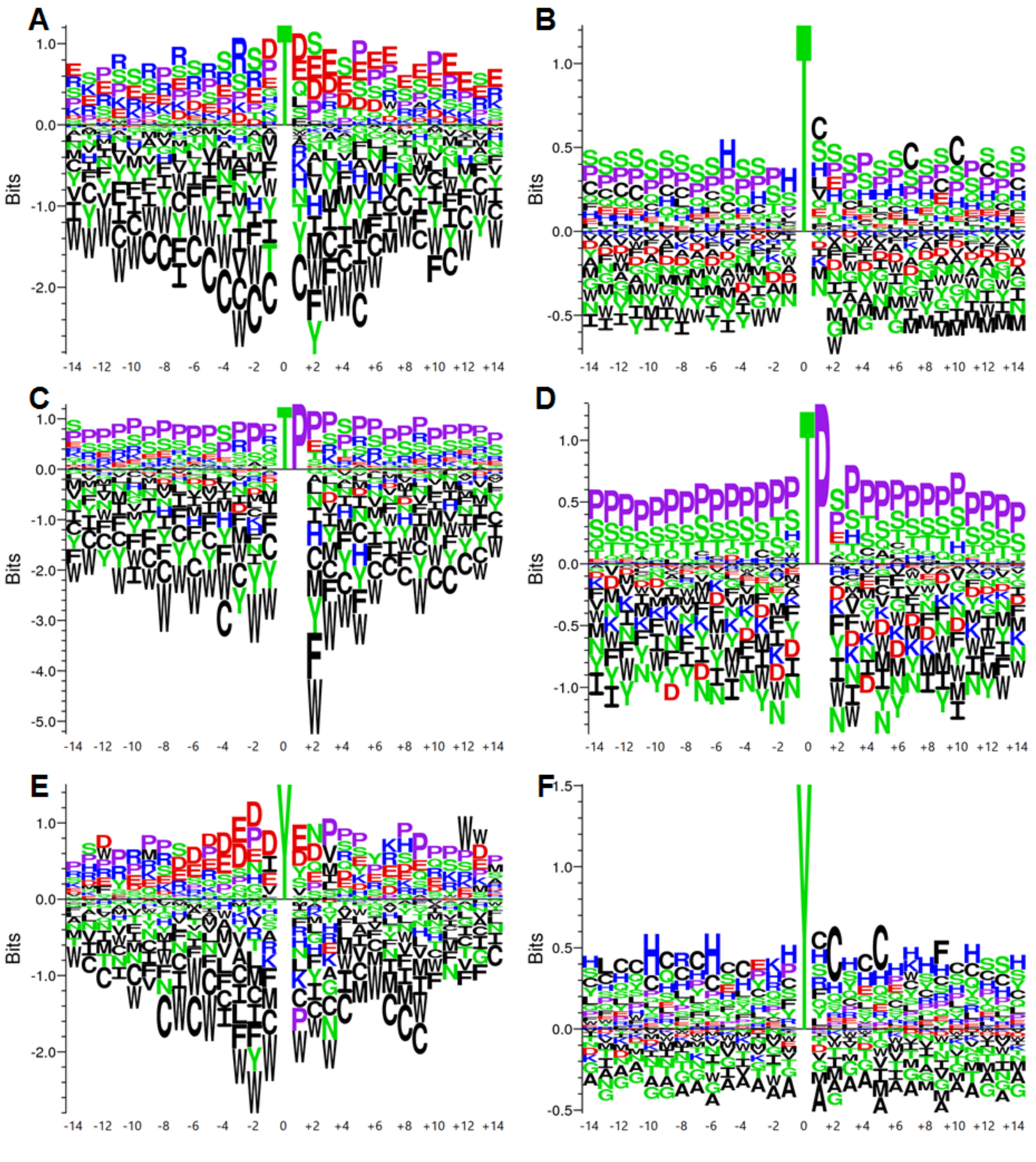


**Fig. S1. Sequence logos of Threonine and Tyrosine phosphorylated or non-phosphorylated amino acid sequence neighborhoods.** A. Phosphorylated Threonine, non-+1 Proline. B. Non-phosphorylated Threonine, non-+1 Proline. C. Phosphorylated Threonine, +1 Proline. D. Non-Phosphorylated Threonine, +1 Proline. E. Phosphorylated Tyrosine. F. Non-Phosphorylated Tyrosine. In all figures, aliphatic/non-polar residues are colored black, prolines are lavender, polar residues are green, negatively charged side chains are red, positively charged side chains are blue.

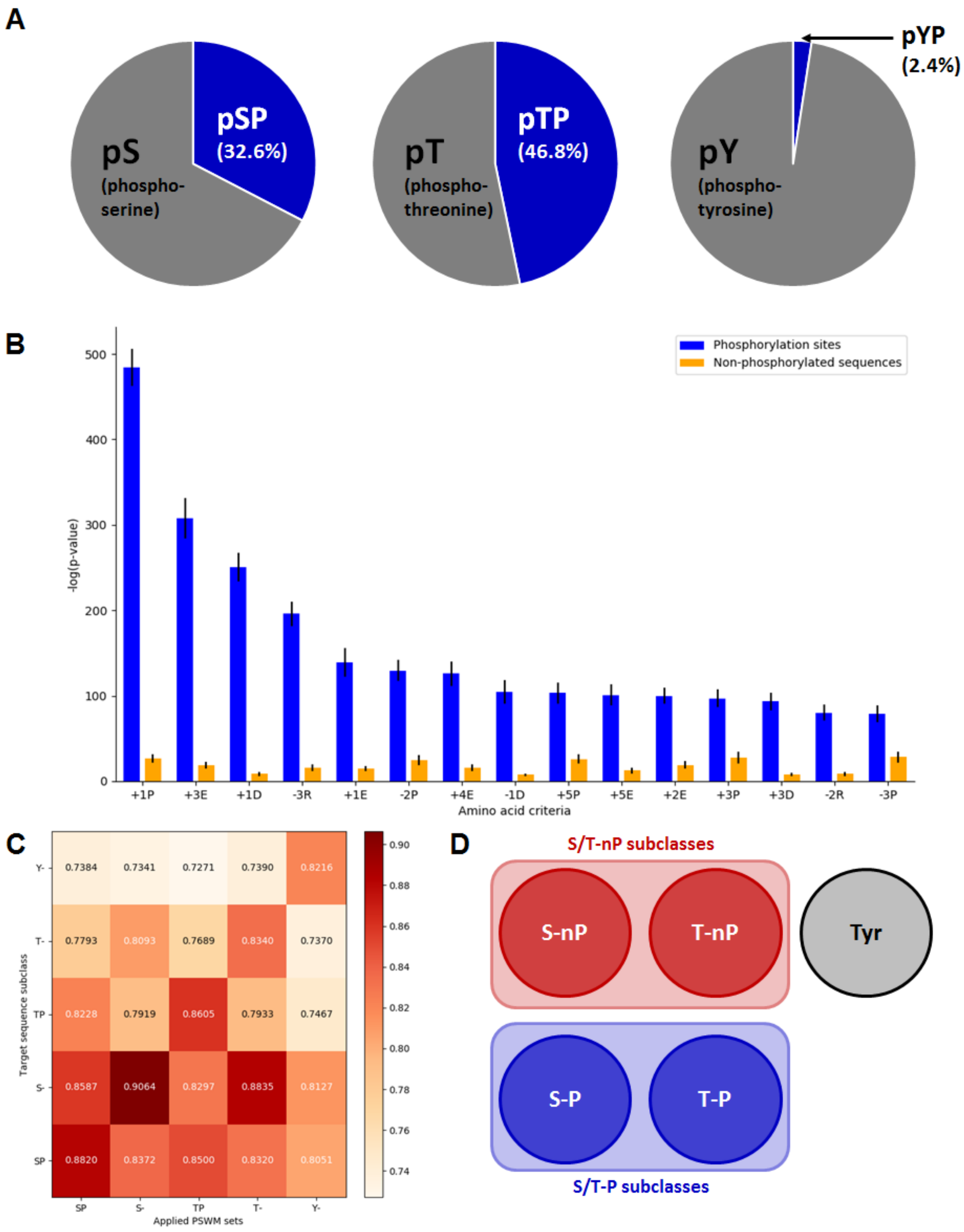


**Fig. S2. Dividing phosphorylation sites by presence or absence of +1 Proline reveals two distinct subclasses in Serine and Theronine phosphorylation sites.** A. Frequencies of +1 Proline phosphorylation sites make up one-third to one-half of the human phosphoproteome. In contrast, Tyrosine phosphorylation sites have few +1 Proline positions. B. Dividing the full set of human Ser/Thr phosphorylation sites into two groups based on the presence or absence of position-specific amino acids reveals that, while many positions contain significant grouping information, the +1 Proline residue is the single most informative; i.e. this single (residue, position) pair results in the most statistically significant subsets. Statistical differences were measured as an average of p-values calculated from t-test conducted for each possible amino acid occurrence at each possible type of site. The four (residue, position) pairs - +1P, +3E, +1D, -3R – which showed the largest average of p-values were selected for each case. Blue bars indicate sequences with known phosphorylation sites, orange bars indicate non-phosphorylated sequences. C. Position-specific weight matrices (PSWM) comparisons between different phosphorylation site subclasses. Prediction of phosphorylation sites with other subclass parameter sets reveal that there is more similarity between classes with the same presence or absence of the +1 Pro residue than between classes with the same type of phosphorylated residue. Scale indicates AUROC value for each prediction result. D. Two possible grouping schemes supported by these analyses are: five subclasses (circles), treating Serine and Threonine as separate subclasses, and three subclasses (rectangles), merging Serine and Threonine subclasses.

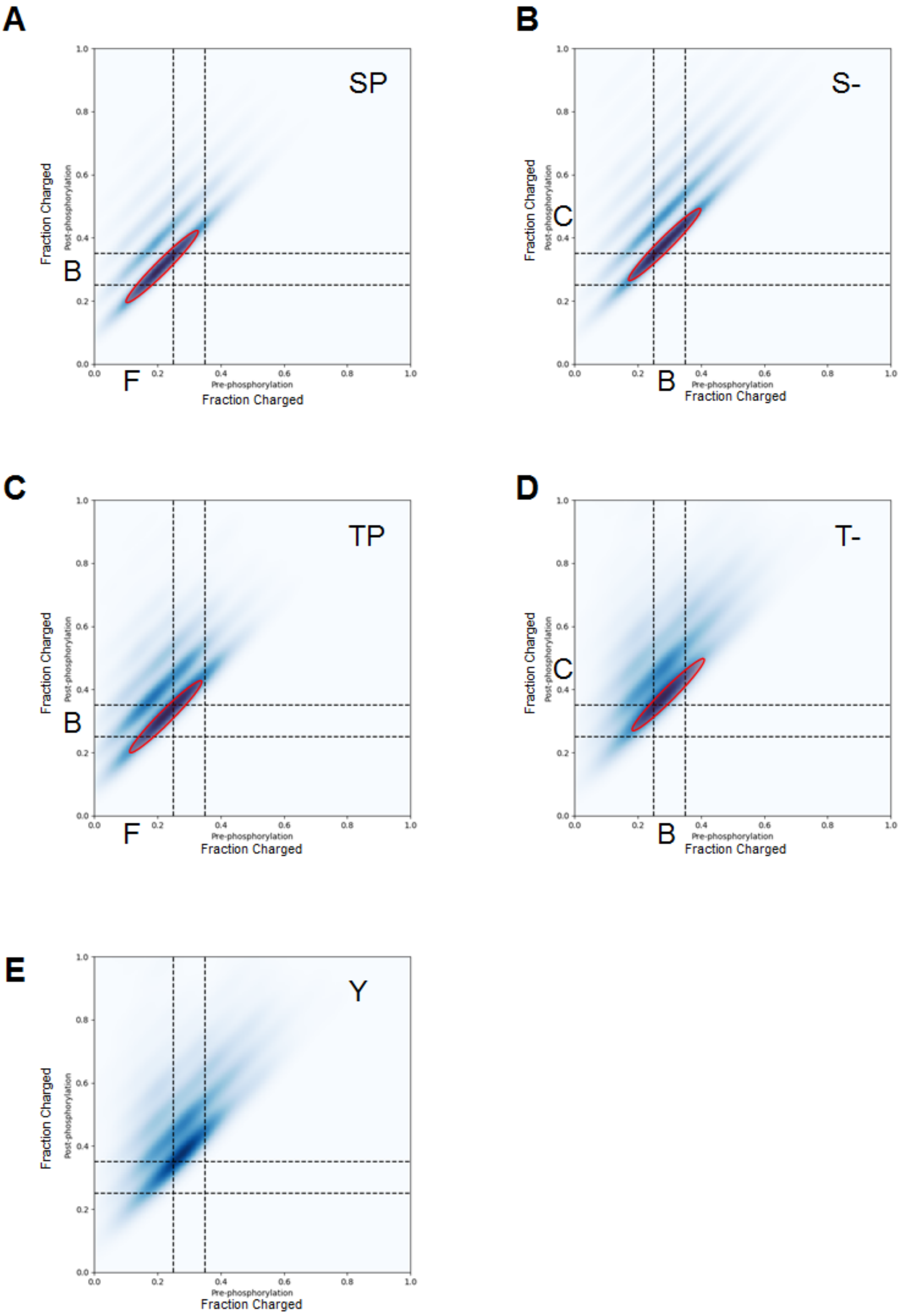
A
SP
Fraction Charged
Post-phosphorylation
Pre-phosphorylation
Fraction Charged
B
F
B
S-
C
B
C
TP
D
T-
E
Y
0.0
0.2
0.4
0.6
0.8
1.0

**Fig. S3. Changes in charge distributions before and after phosphorylation events.** A. Ser +1 Pro sites. B. Ser non +1 Pro sites. C. Thr +1 Pro sites. D. Thr non +1 Pro sites. E. Tyr sites. In each panel, individual points represent one 29-mer sequence, with average charge fractions plotted before (x-axis) or after (y-axis) a single, double, triple, or quadruple phosphorylation events adding successive negative charges. Dashed lines indicate boundary regions defined by Das & Pappu (15): B=boundary region 2, F=folded region 1, C=coil regions 3,4,5. Red circles emphasize the differential shifts in the conformational manifold of each subclass upon a single phosphorylation event as described in detail in Main Text Figures 4C-D.

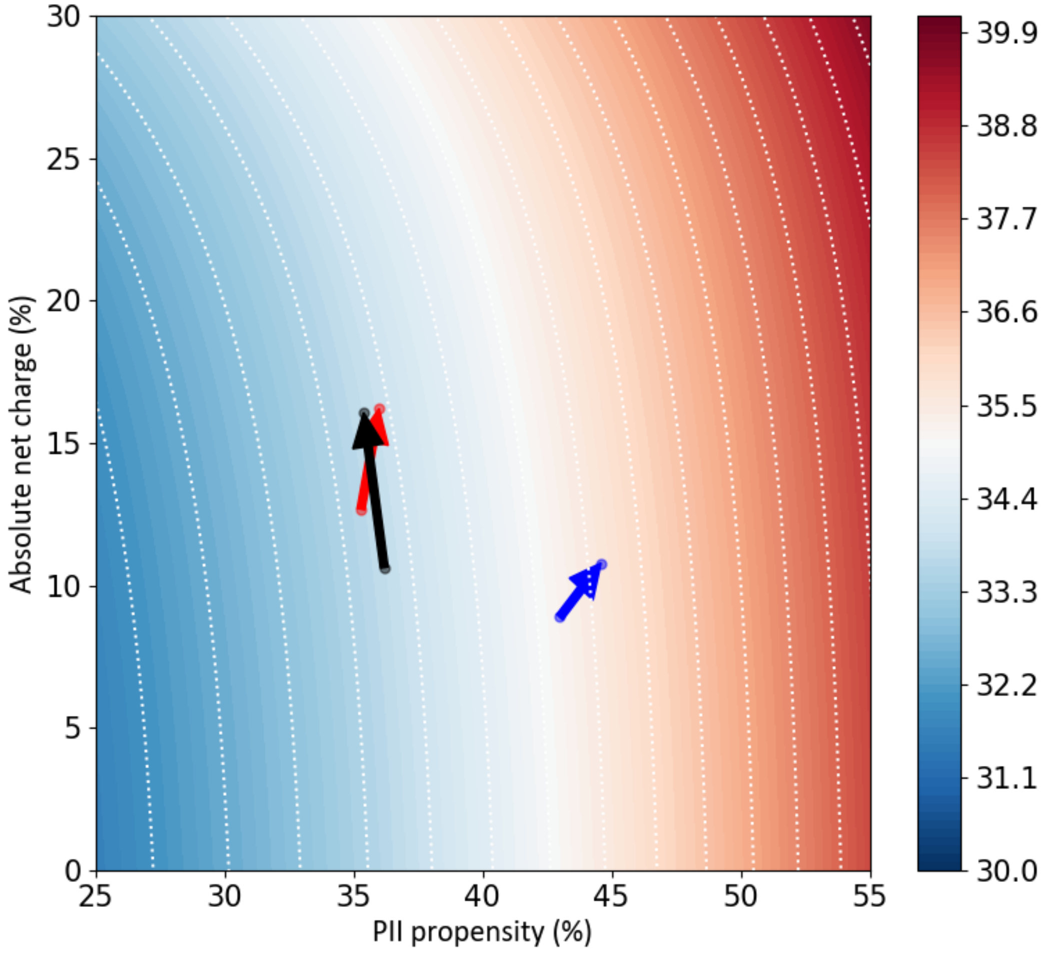


**Fig. S4. Phosphorylation sites containing +1 Proline are energetically poised to respond to phosphorylation by extension, mediated by charge and polyproline II propensity.** Red, white, and blue contour regions indicate predicted end-to-end distances of intrinsically disordered proteins from a theoretical model (16) that takes polyproline II structural propensity (x-axis) and net charge (y-axis) into account. Arrows on this contour plot indicate median predicted distances of distributions of known phosphorylation sites before (arrow tail) and after (arrow head) a single phosphorylation event. Red arrow denotes Serine/Threonine non +1 Proline sites, blue arrow denotes Serine/Threonine +1 Proline sites, and Black arrow denotes Tyrosine sites. Scale bar indicates end-to-end distance in Ångstroms.

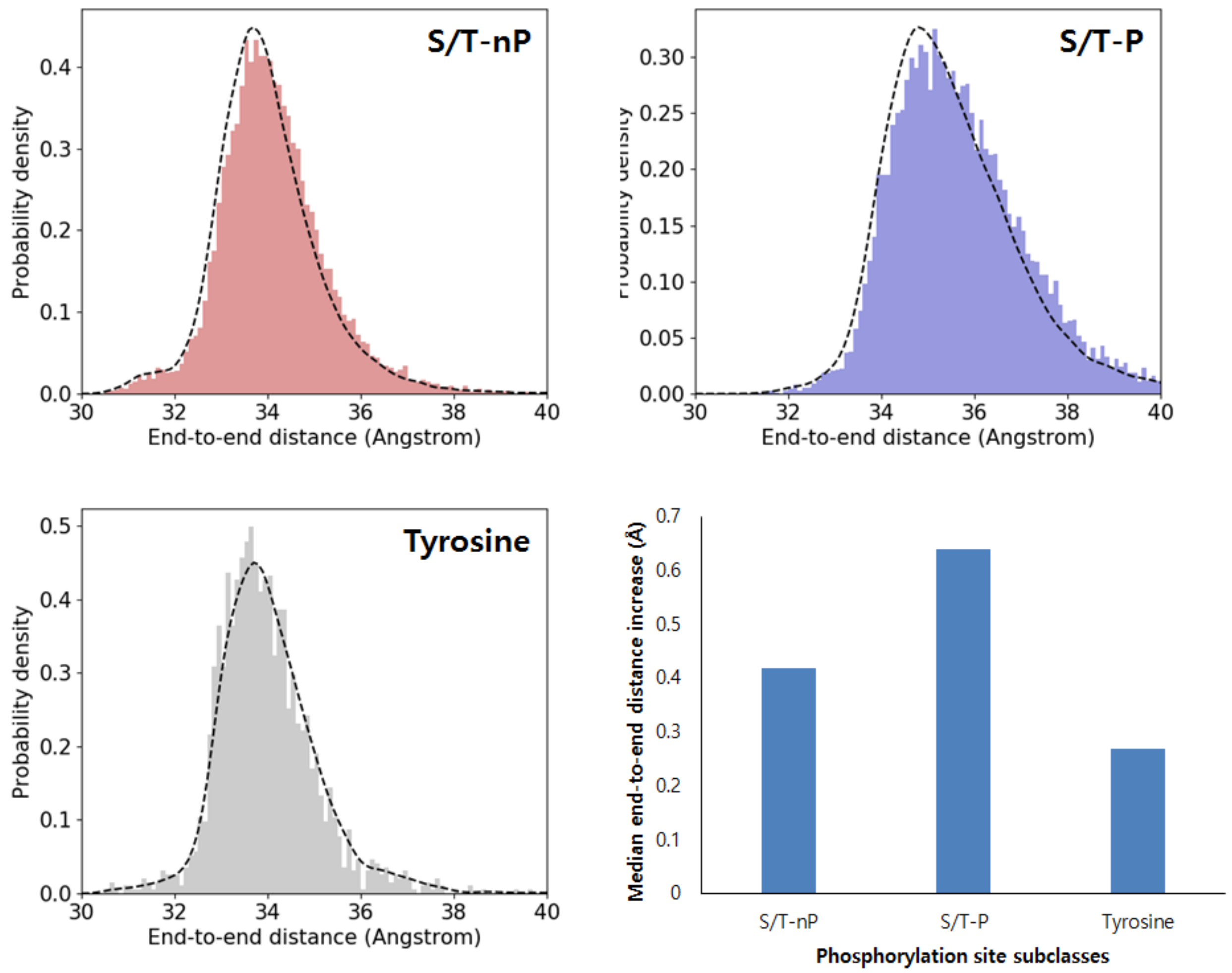


**Fig. S5. Phosphorylation site subclasses defined with +1 Proline show higher end-to-end distances than other subclasses.** In these distributions, red represents sequences before phosphorylation, blue represents sequences after phosphorylation, and purple represents areas of overlap between red and blue; a smaller area of overlap thus suggests a greater change of end-to-end distance after phosphorylation. The column plot demonstrates the median distance increase for each case, with the +1 Proline sites exhibiting the largest increase.

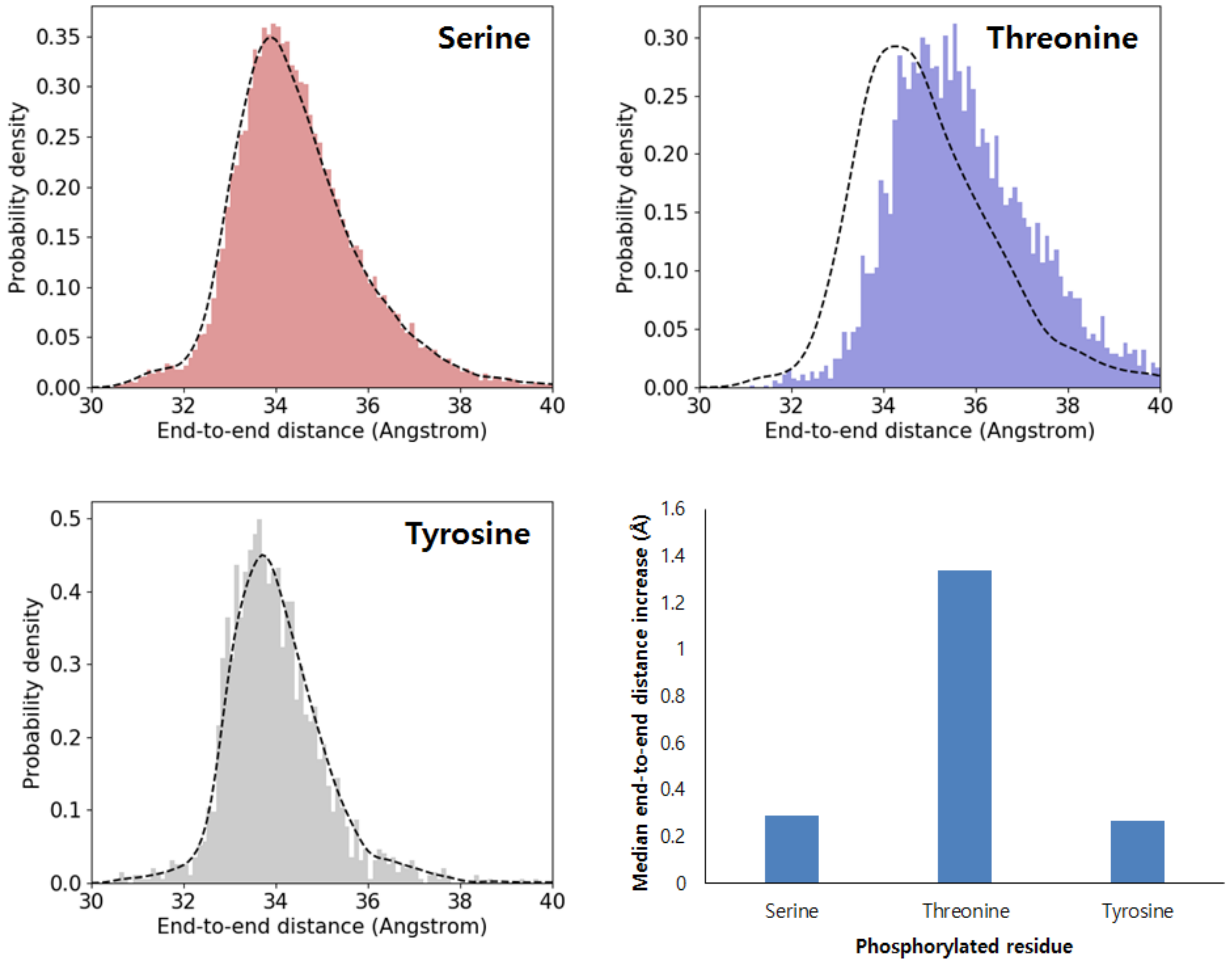


**Fig. S6. Threonine phosphorylation has a stronger effect on end-to-end distance increase than do Serine / Tyrosine phosphorylations.** In these distributions, red represents sequences before phosphorylation, blue represents sequences after phosphorylation, and purple represents areas of overlap between red and blue; a smaller area of overlap thus suggests a greater change of end-to-end distance after phosphorylation. The column plot demonstrates the median distance increase for each case, with the Threonine sites exhibiting the largest increase, more than one Å after phosphorylation.

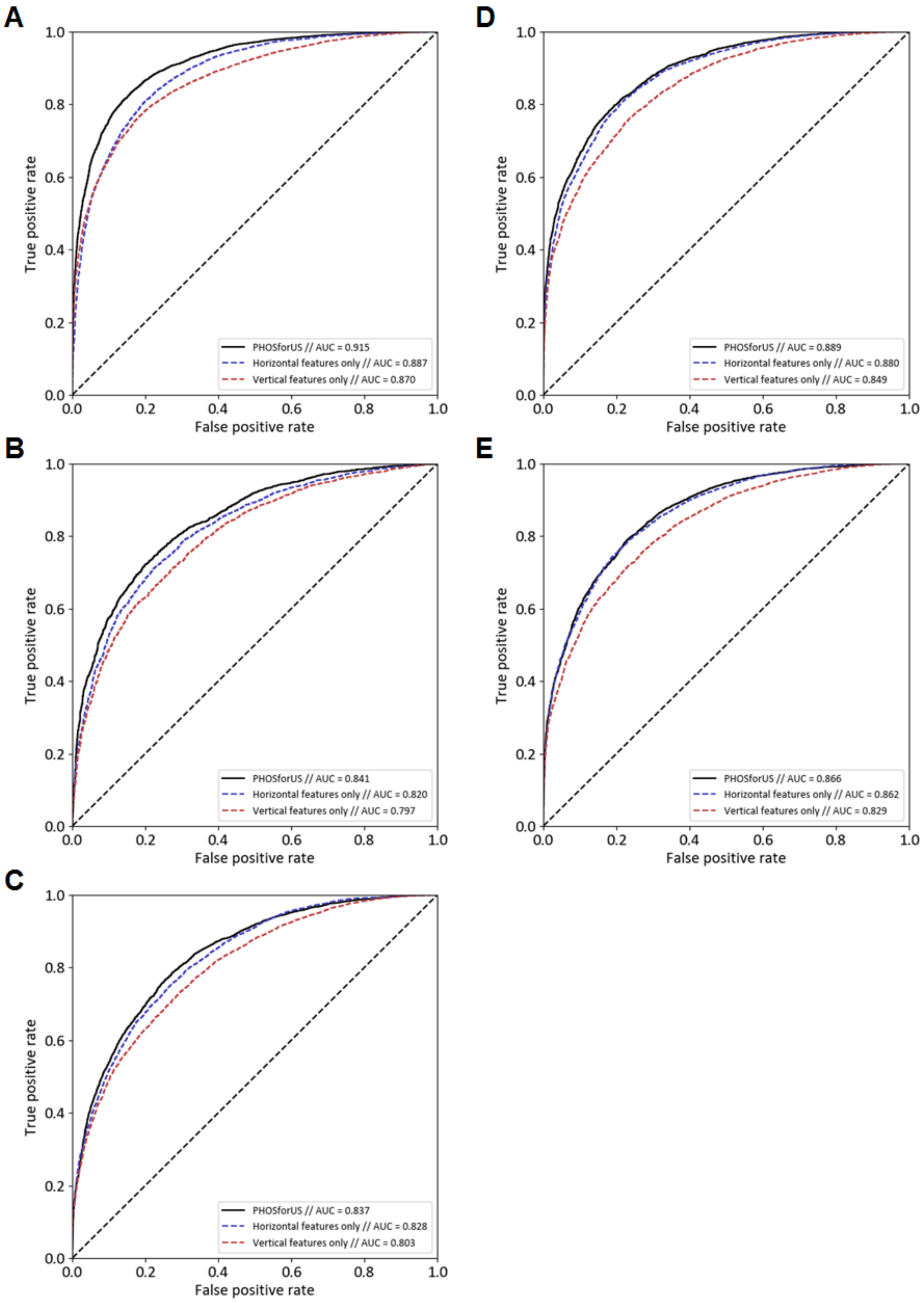
A
B
C
D
E
True positive rate
False positive rate
PHOSforUS // AUC = 0.915
Horizontal features only // AUC = 0.887
Vertical features only // AUC = 0.870
PHOSforUS // AUC = 0.841
Horizontal features only // AUC = 0.820
Vertical features only // AUC = 0.797
PHOSforUS // AUC = 0.837
Horizontal features only // AUC = 0.828
Vertical features only // AUC = 0.803
PHOSforUS // AUC = 0.889
Horizontal features only // AUC = 0.880
Vertical features only // AUC = 0.849
PHOSforUS // AUC = 0.866
Horizontal features only // AUC = 0.862
Vertical features only // AUC = 0.829

**Fig. S7. Subclass-specific receiver–operating characteristics (ROC) of PHOSforUS constituent predictors.** AUROC stands for “Area Under the ROC curve”. For all subclasses, predictors using horizontal information are equivalent to, or more effective than, predictors using vertical information. A. Serine non-+1 Proline sites. B. Threonine non-+1 proline sites. C. Tyrosine sites. D. Serine +1 Proline sites. E. Threonine +1 proline sites.

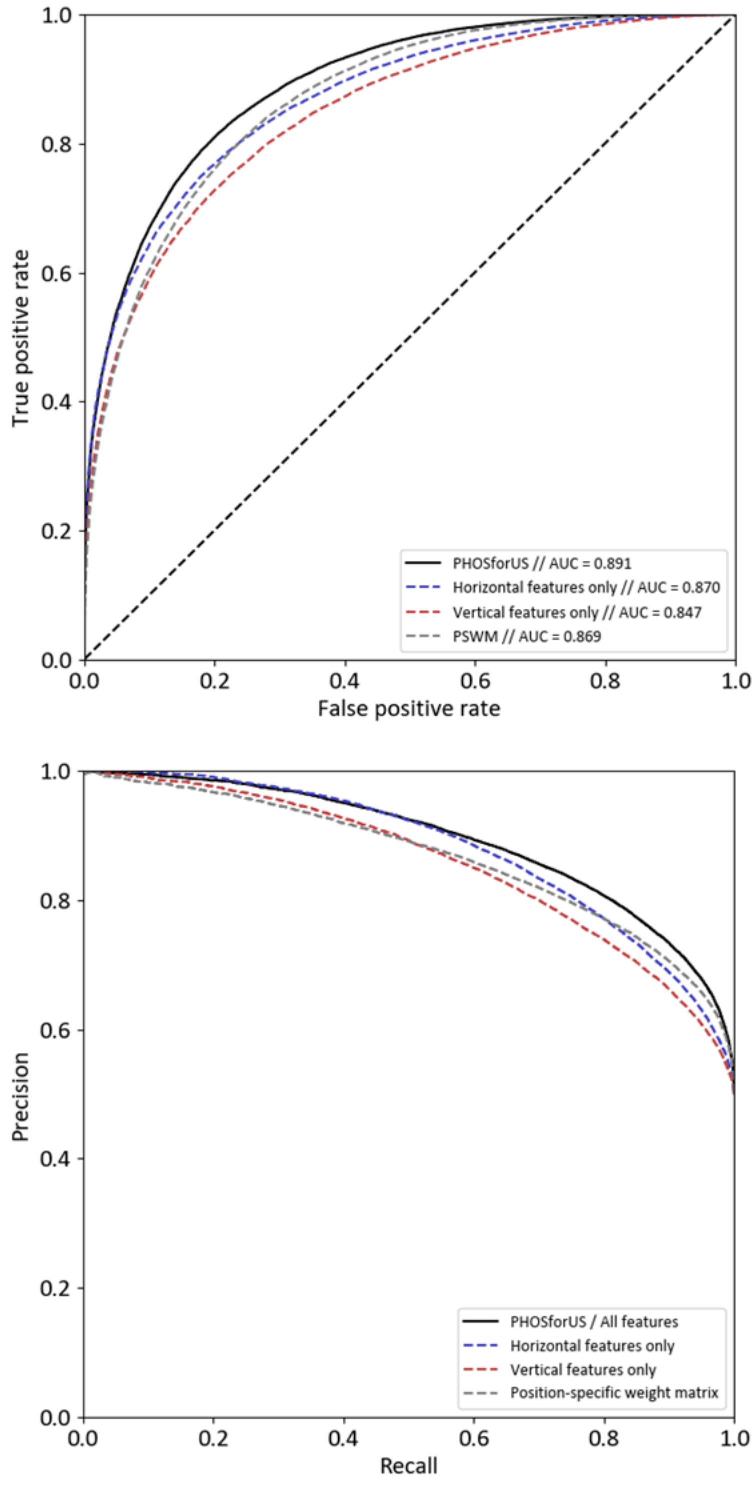


**Fig. S8. Receiver-operating characteristics (ROC) curve (upper panel) and precision-recall curve (lower panel) of PHOSforUS predictor & its subpredictors along with PSWM-based prediction results.** These results are based on the same data displayed in Figures 5B-D of the main text and Supplementary Figure S9A, below.

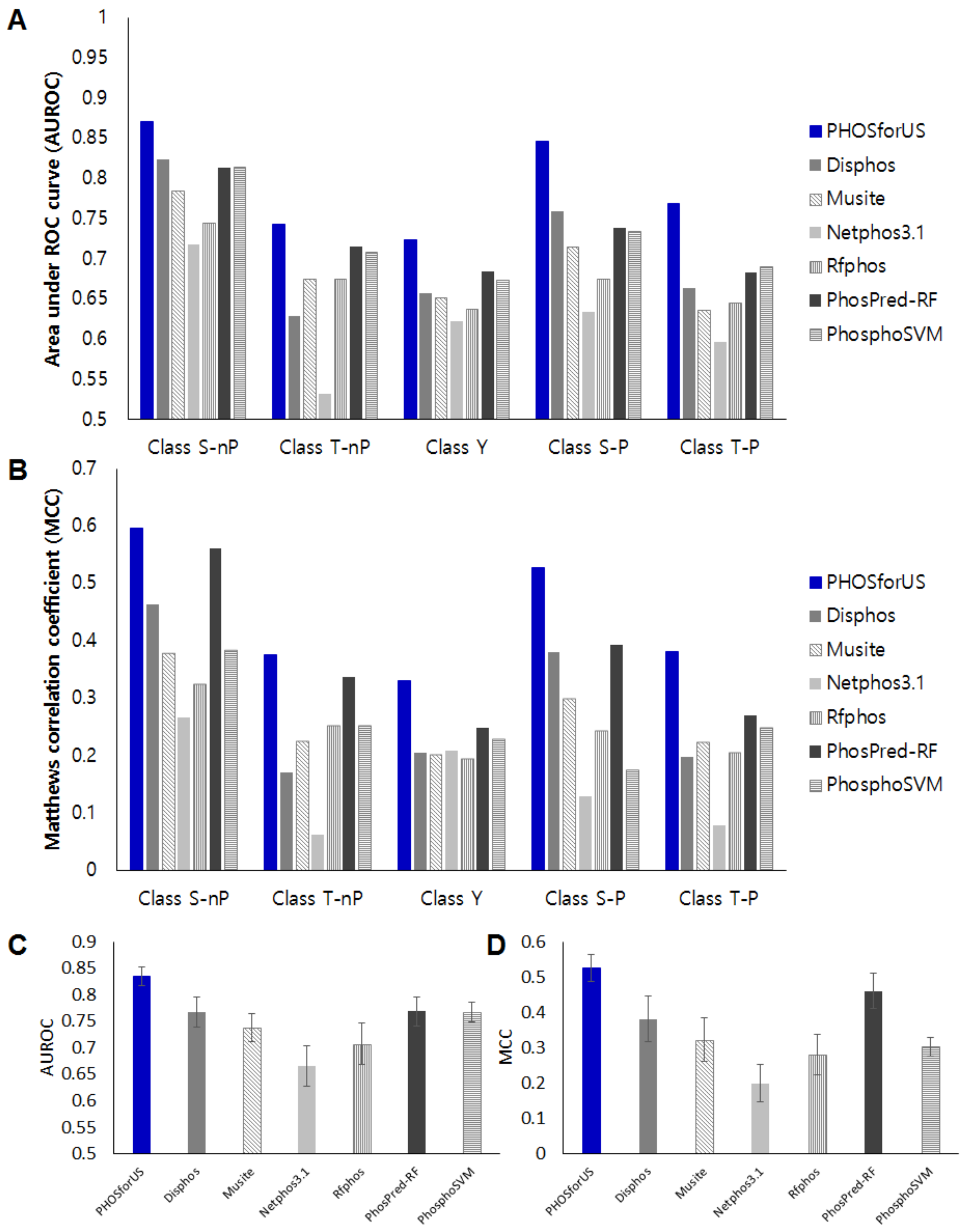


**Fig. S9. Comparative effectiveness of protein phosphorylation site prediction by PHOSforUS**. A. Class-specific AUROC, displayed again in Figures 5B-D of the Main Text. B. Class-specific Matthews Correlation Coefficient (MCC). C. Weighted average of AUROC. D. Weighted average of MCC

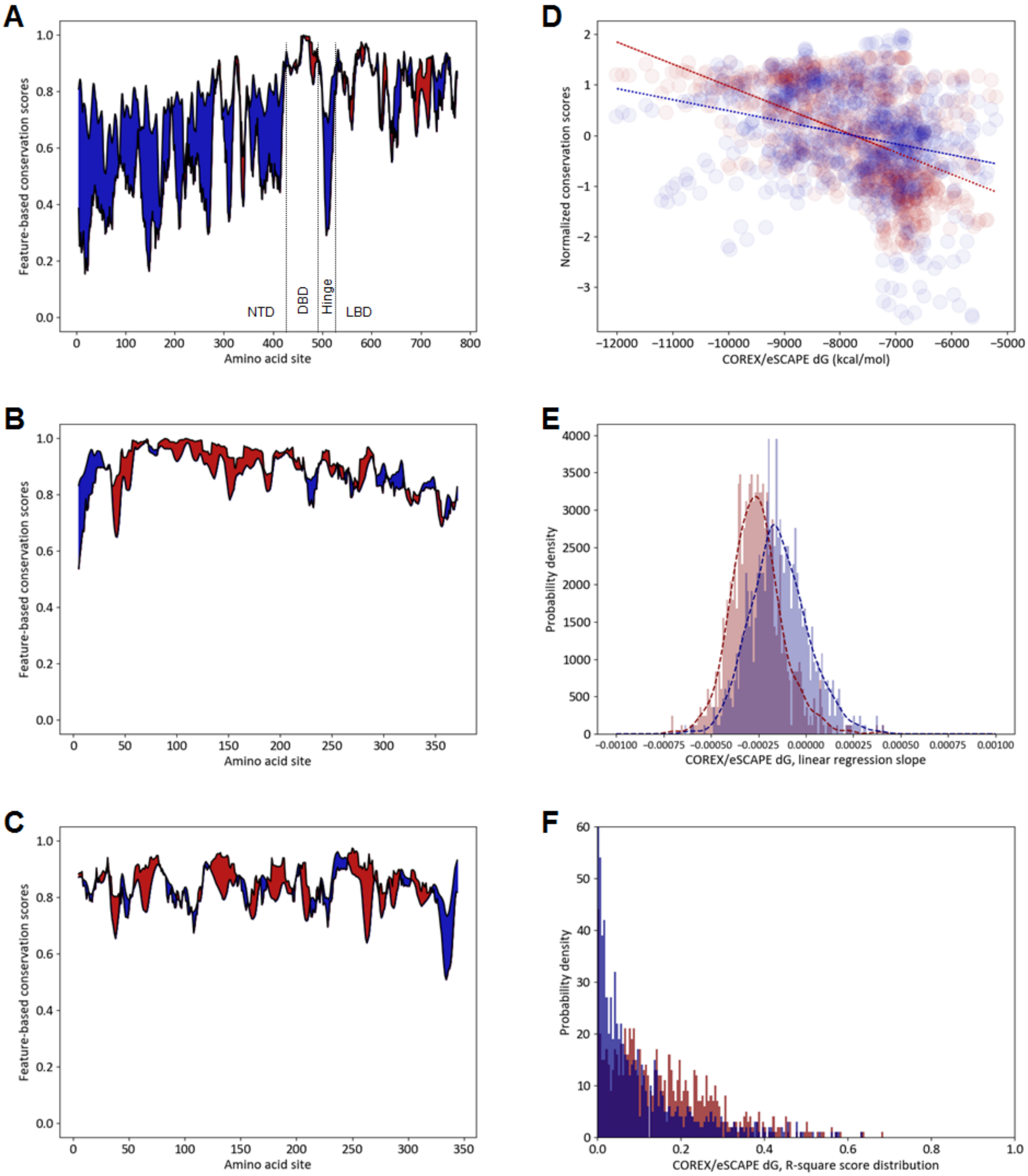


**Fig. S10. Horizontal information is better conserved than vertical information in intrinsically disordered region.** In all panels, red indicates conservation of vertical information and blue indicates conservation of horizontal information. A. Difference between degrees of conservation of sequence and free energy (ΔG, (5)) calculated for human glucocorticoid receptor (OMA database identifier GR) and its orthologs (17). Free energy is used as an example of horizontal information, and amino acid sequence conservation is used as an example of vertical information. Conservation is computed as described above in Methods and is normalized using Equation S11, above. Blue denotes regions where free energy conservation is stronger than sequence conservation, and red denotes the opposite. In human GR, DNA binding region (DBD) and LBD region are structured, while N-terminal domain (NTD) and hinge region are intrinsically disordered. B. Same calculation for actin (OMA database identifier ACTB). C. Same calculation for rhodopsin (OMA database identifier RHO). D. Correlation between COREX/eSCAPE ΔG and normalized conservation scores for individual residue positions in the GR family. Red: sequence conservation score, Blue: ΔG conservation score. The shallower slope of the blue line suggests that ΔG conservation is stronger than amino acid sequence conservation for this family. E. Distribution of linear regression slopes for 835 different transcription families. The median slope of the blue distribution being closer to zero suggests that ΔG conservation is stronger than amino acid conservation for this large collection of different protein families. Thus, strong conservation of horizontal information seems to be a general property of protein evolution. Red: sequence conservation, Blue: ΔG conservation. F. Distribution of R-square values of linear regression. Lower median R2 values for ΔG conservation also suggests that horizontal information is more strongly conserved than vertical information in this large collection of different protein families.

**Supplementary Tables**

**Table S1. Statistics of utilized sequence & annotation datasets.**

| | Total P-sites | | | Total N-sites | |
|---|---|---|---|---|---|
| **Class** | **Pre-screening** | **After screening** | **Comparative analysis** | **Pre-screening** | **After screening** |
| **S-P** | 10348 | 3024 | 11842 | 30170 | 7373 |
| **S-nP** | 21936 | 4426 | 55628 | 455303 | 88905 |
| **T-P** | 2688 | 1176 | 5028 | 20943 | 1762 |
| **T-nP** | 3045 | 1385 | 20299 | 288492 | 27627 |
| **Y** | 2058 | 1145 | 14415 | 145170 | 24271 |

**Table S2. List of biophysical indices incorporated in PHOSforUS predictor.**

| Feature ID | Description | Feature type | Reference |
|---|---|---|---|
| GUYH850101† | Partition energy | Hydrophobicity / Horizontal | Guy (1985) |
| MIYS990104† | Optimized relative partition energy - method C | Hydrophobicity / Horizontal | Miyazawa-Jernigan (1994) |
| PRAM900102† | Relative frequency in alpha-helix | Conformation / Horizontal | Prabhakaran (1990) |
| PALJ810112† | Normalized frequency of beta-sheet | Conformation / Horizontal | Palau et al. (1981) |
| ROBB760105† | Information measure for extended | Conformation / Horizontal | Robson-Suzuki (1976) |
| PPIIPRO | Polyproline II propensity | Conformation / Horizontal | Elam et al. (2013) (3) |
| ZIMJ680104† | Isoelectric points | Vertical | Zimmerman et al. (1968) |
| FASG760101† | Molecular weight | Vertical | Fasman (1976) |
| GRAR740103† | Residue volume | Vertical | Grantham (1974) |
| RADA880106† | Accessible surface area | Vertical | Radzicka-Wolfenden (1988) |

† Feature IDs correspond to the scales contained in AAindex (1)

**Table S3. List of eSCAPE thermodynamic parameters incorporated in PHOSforUS predictor.**

| ***eSCAPE* parameter** | **Description** |
|---|---|
| $\Delta G$,N | Gibbs free energy of folded state |
| $\Delta H_{ap}$,N | Apolar enthalpy of folded state |
| $\Delta H_{pol}$,N | Polar enthalpy of folded state |
| $T\Delta S_{conf}$,N | Conformational entropy of folded state |
| $\Delta\Delta G$ ($\Delta G$,N – $\Delta G$,D) | $\Delta G$ difference between folded & unfolded state |
| $\Delta\Delta H_{ap}$ ($\Delta H_{ap}$,N – $\Delta H_{ap}$,D) | $\Delta H_{ap}$ difference between folded & unfolded state |
| $\Delta\Delta H_{pol}$ ($\Delta H_{pol}$,N – $\Delta H_{pol}$,D) | $\Delta H_{pol}$ difference between folded & unfolded state |
| $\Delta T\Delta S_{conf}$ ($T\Delta S$,N – $T\Delta S$,D) | $T\Delta S_{conf}$ difference between folded & unfolded state |

**Table S4. Sub-predictor statistics for Serine +1 Proline (S-P) subclass.** Values in red font indicate the largest statistic value in each feature group.

| **Class S-P** | **Accuracy** | **Sensitivity** | **Specificity** | **Precision** | **F1** | **MCC** | **AUROC** |
|---|---|---|---|---|---|---|---|
| ZIMJ680104 | 0.598934 | 0.549289 | 0.648578 | 0.609838 | 0.577963 | 0.198858 | 0.640449 |
| FASG760101 | 0.644826 | 0.646998 | 0.642654 | 0.644284 | 0.645597 | 0.289692 | 0.698131 |
| GRAR740103 | 0.680016 | 0.664929 | 0.695103 | 0.685712 | 0.675114 | 0.360244 | 0.743468 |
| RADA880106 | 0.673697 | 0.620458 | 0.726935 | 0.6945 | 0.655319 | 0.34945 | 0.740499 |
| **Vertical features** | **0.752725** | **0.738784** | **0.766667** | **0.760029** | **0.749216** | **0.505698** | **0.835519** |
| GUYH850101 | 0.752765 | 0.76722 | 0.73831 | 0.745754 | 0.756271 | 0.505829 | 0.834035 |
| MIYS990104 | 0.759874 | 0.779226 | 0.740521 | 0.750265 | 0.76444 | 0.52018 | 0.840712 |
| PRAM900102 | 0.619471 | 0.517615 | 0.721327 | 0.649957 | 0.576162 | 0.244081 | 0.663511 |
| PALJ810112 | 0.664218 | 0.704265 | 0.624171 | 0.652126 | 0.677157 | 0.329528 | 0.724739 |
| ROBB760105 | 0.702291 | 0.732148 | 0.672433 | 0.690894 | 0.710886 | 0.405357 | 0.774226 |
| PPIIPRO | 0.662046 | 0.530174 | 0.793918 | 0.720275 | 0.610634 | 0.336092 | 0.724 |
| $\Delta G$,N | 0.683965 | 0.709795 | 0.658136 | 0.674974 | 0.691914 | 0.368459 | 0.748253 |
| $\Delta H_{ap}$,N | 0.612243 | 0.539652 | 0.684834 | 0.631285 | 0.581816 | 0.226914 | 0.657694 |
| $\Delta H_{pol}$,N | 0.622749 | 0.674724 | 0.570774 | 0.611197 | 0.641368 | 0.246861 | 0.667824 |
| $T\Delta S_{conf}$,N | 0.606635 | 0.671248 | 0.542022 | 0.594464 | 0.630492 | 0.215103 | 0.642729 |
| $\Delta\Delta G$,N-D | 0.693009 | 0.65158 | 0.734439 | 0.710504 | 0.679719 | 0.387398 | 0.762268 |
| $\Delta\Delta H_{ap}$,N-D | 0.674171 | 0.71169 | 0.636651 | 0.662028 | 0.685947 | 0.349344 | 0.736206 |
| $\Delta\Delta H_{pol}$,N-D | 0.635427 | 0.590758 | 0.680095 | 0.648688 | 0.618317 | 0.271969 | 0.687678 |
| $T\Delta\Delta S_{conf}$,N-D | 0.667457 | 0.627409 | 0.707504 | 0.682081 | 0.653554 | 0.336035 | 0.728165 |
| **Horizontal features** | **0.78207** | **0.792733** | **0.771406** | **0.77631** | **0.784389** | **0.564335** | **0.870907** |
| **Total features** | **0.794589** | **0.800158** | **0.789021** | **0.791433** | **0.795736** | **0.589268** | **0.882882** |

**Table S5. Sub-predictor statistics for Serine non +1 Proline (S-nP) subclass.** Values in red font indicate the largest statistic value in each feature group.

| **Class S-nP** | **Accuracy** | **Sensitivity** | **Specificity** | **Precision** | **F1** | **MCC** | **AUROC** |
|---|---|---|---|---|---|---|---|
| ZIMJ680104 | 0.697775 | 0.67005 | 0.7255 | 0.709437 | 0.689139 | 0.396203 | 0.762915 |
| FASG760101 | 0.6173 | 0.63695 | 0.59765 | 0.612898 | 0.62467 | 0.234798 | 0.669534 |
| GRAR740103 | 0.678525 | 0.6728 | 0.68425 | 0.68063 | 0.676664 | 0.357103 | 0.744073 |
| RADA880106 | 0.6253 | 0.58055 | 0.67005 | 0.6376 | 0.607705 | 0.251627 | 0.672302 |
| **Vertical features** | **0.791125** | **0.7627** | **0.81955** | **0.808718** | **0.785009** | **0.583229** | **0.873915** |
| GUYH850101 | 0.78475 | 0.8062 | 0.7633 | 0.773094 | 0.789258 | 0.570093 | 0.867598 |
| MIYS990104 | 0.790275 | 0.8058 | 0.77475 | 0.781566 | 0.793458 | 0.580896 | 0.871936 |
| PRAM900102 | 0.58555 | 0.46655 | 0.70455 | 0.612254 | 0.529459 | 0.176183 | 0.618341 |
| PALJ810112 | 0.709825 | 0.72305 | 0.6966 | 0.704479 | 0.71362 | 0.419824 | 0.786046 |
| ROBB760105 | 0.7535 | 0.75005 | 0.75695 | 0.755266 | 0.752613 | 0.507059 | 0.838519 |
| PPIIPRO | 0.597175 | 0.42545 | 0.7689 | 0.647984 | 0.513567 | 0.206941 | 0.632913 |
| $\Delta G$,N | 0.695025 | 0.7278 | 0.66225 | 0.683054 | 0.70469 | 0.390927 | 0.769178 |
| $\Delta H_{ap}$,N | 0.546275 | 0.6325 | 0.46005 | 0.539487 | 0.582283 | 0.093961 | 0.57366 |
| $\Delta H_{pol}$,N | 0.617125 | 0.69 | 0.54425 | 0.602246 | 0.643121 | 0.236801 | 0.661454 |
| $T\Delta S_{conf}$,N | 0.652675 | 0.6847 | 0.62065 | 0.643589 | 0.663469 | 0.306011 | 0.708972 |
| $\Delta\Delta G$,N-D | 0.66655 | 0.67195 | 0.66115 | 0.664795 | 0.668329 | 0.333145 | 0.727326 |
| $\Delta\Delta H_{ap}$,N-D | 0.69165 | 0.73315 | 0.65015 | 0.677023 | 0.703932 | 0.384673 | 0.764952 |
| $\Delta\Delta H_{pol}$,N-D | 0.600525 | 0.6195 | 0.58155 | 0.596944 | 0.607967 | 0.201224 | 0.638616 |
| $T\Delta\Delta S_{conf}$,N-D | 0.647875 | 0.6513 | 0.64445 | 0.646919 | 0.649078 | 0.295779 | 0.704685 |
| **Horizontal features** | **0.818325** | **0.82745** | **0.8092** | **0.81277** | **0.819977** | **0.636863** | **0.898609** |
| **Total features** | **0.8376** | **0.8432** | **0.832** | **0.833922** | **0.838486** | **0.675324** | **0.918708** |

**Table S6. Sub-predictor statistics for Threonine +1 Proline (T-P) subclass.** Values in red font indicate the largest statistic value in each feature group.

| Class T-P | Accuracy | Sensitivity | Specificity | Precision | F1 | MCC | AUROC |
|---|---|---|---|---|---|---|---|
| ZIMJ680104 | 0.596562 | 0.551003 | 0.64212 | 0.60613 | 0.577107 | 0.193978 | 0.635436 |
| FASG760101 | 0.608596 | 0.624642 | 0.59255 | 0.605623 | 0.614697 | 0.2175 | 0.660287 |
| GRAR740103 | 0.634241 | 0.626074 | 0.642407 | 0.63686 | 0.63117 | 0.268733 | 0.682833 |
| RADA880106 | 0.641404 | 0.59341 | 0.689398 | 0.656395 | 0.62315 | 0.28421 | 0.687897 |
| **Vertical features** | **0.703295** | **0.709742** | **0.696848** | **0.700794** | **0.705197** | **0.406673** | **0.779973** |
| GUYH850101 | 0.691977 | 0.735817 | 0.648138 | 0.67625 | 0.704671 | 0.385647 | 0.766355 |
| MIYS990104 | 0.704155 | 0.743266 | 0.665043 | 0.689275 | 0.715105 | 0.409807 | 0.775734 |
| PRAM900102 | 0.600573 | 0.497708 | 0.703438 | 0.626777 | 0.554658 | 0.205645 | 0.631605 |
| PALJ810112 | 0.632378 | 0.689685 | 0.575072 | 0.619099 | 0.652177 | 0.266828 | 0.684848 |
| ROBB760105 | 0.659456 | 0.712034 | 0.606877 | 0.64435 | 0.676375 | 0.320855 | 0.711871 |
| PPIIPRO | 0.640258 | 0.513467 | 0.767049 | 0.687936 | 0.587811 | 0.290073 | 0.702787 |
| $\Delta G$,N | 0.637536 | 0.667622 | 0.60745 | 0.629882 | 0.647905 | 0.275879 | 0.695882 |
| $\Delta H_{ap}$,N | 0.582665 | 0.470774 | 0.694556 | 0.606579 | 0.529857 | 0.169718 | 0.613442 |
| $\Delta H_{pol}$,N | 0.603295 | 0.658453 | 0.548138 | 0.593015 | 0.623854 | 0.208042 | 0.643289 |
| $T\Delta S_{conf}$,N | 0.574355 | 0.659026 | 0.489685 | 0.563547 | 0.607445 | 0.151014 | 0.60039 |
| $\Delta\Delta G$,N-D | 0.64298 | 0.607163 | 0.678797 | 0.653712 | 0.629321 | 0.286842 | 0.701574 |
| $\Delta\Delta H_{ap}$,N-D | 0.620917 | 0.661032 | 0.580802 | 0.612181 | 0.635283 | 0.242993 | 0.67731 |
| $\Delta\Delta H_{pol}$,N-D | 0.602579 | 0.537249 | 0.667908 | 0.61822 | 0.574768 | 0.207032 | 0.649613 |
| $T\Delta\Delta S_{conf}$,N-D | 0.633954 | 0.575358 | 0.69255 | 0.651677 | 0.610863 | 0.269926 | 0.684493 |
| **Horizontal features** | **0.723782** | **0.760172** | **0.687393** | **0.708504** | **0.733173** | **0.449166** | **0.80064** |
| **Total features** | **0.741404** | **0.767908** | **0.7149** | **0.729282** | **0.747969** | **0.483678** | **0.8199** |

**Table S7. Sub-predictor statistics for Threonine non +1 Proline (T-nP) subclass.** Values in red font indicate the largest statistic value in each feature group.

| Class T-nP | Accuracy | Sensitivity | Specificity | Precision | F1 | MCC | AUROC |
|---|---|---|---|---|---|---|---|
| ZIMJ680104 | 0.617098 | 0.583364 | 0.650832 | 0.625888 | 0.60373 | 0.234852 | 0.660006 |
| FASG760101 | 0.57403 | 0.609057 | 0.539002 | 0.569071 | 0.588254 | 0.148527 | 0.602642 |
| GRAR740103 | 0.619131 | 0.607763 | 0.630499 | 0.621895 | 0.61446 | 0.23854 | 0.663905 |
| RADA880106 | 0.586784 | 0.521442 | 0.652126 | 0.600085 | 0.557798 | 0.175196 | 0.618641 |
| **Vertical features** | **0.697782** | **0.686322** | **0.709242** | **0.702684** | **0.694134** | **0.39596** | **0.767477** |
| GUYH850101 | 0.699723 | 0.729575 | 0.669871 | 0.688637 | 0.708367 | 0.400345 | 0.770304 |
| MIYS990104 | 0.703974 | 0.721996 | 0.685952 | 0.69702 | 0.709065 | 0.408498 | 0.777033 |
| PRAM900102 | 0.563863 | 0.431608 | 0.696118 | 0.586981 | 0.497185 | 0.132533 | 0.58646 |
| PALJ810112 | 0.646026 | 0.655638 | 0.636414 | 0.643223 | 0.649048 | 0.292422 | 0.70165 |
| ROBB760105 | 0.67366 | 0.674492 | 0.672828 | 0.673373 | 0.673683 | 0.347582 | 0.737826 |
| PPIIPRO | 0.584196 | 0.402403 | 0.765989 | 0.632722 | 0.491213 | 0.181002 | 0.622557 |
| $\Delta G$,N | 0.62597 | 0.653974 | 0.597967 | 0.619301 | 0.635956 | 0.252567 | 0.673932 |
| $\Delta H_{ap}$,N | 0.522089 | 0.479667 | 0.56451 | 0.524836 | 0.500084 | 0.044667 | 0.536126 |
| $\Delta H_{pol}$,N | 0.576617 | 0.61756 | 0.535675 | 0.570771 | 0.593159 | 0.15382 | 0.60663 |
| $T\Delta S_{conf}$,N | 0.597135 | 0.642514 | 0.551756 | 0.589062 | 0.614575 | 0.195121 | 0.642398 |
| $\Delta\Delta G$,N-D | 0.6122 | 0.573937 | 0.650462 | 0.621622 | 0.596631 | 0.225185 | 0.651545 |
| $\Delta\Delta H_{ap}$,N-D | 0.619501 | 0.657671 | 0.581331 | 0.610992 | 0.633263 | 0.239947 | 0.675139 |
| $\Delta\Delta H_{pol}$,N-D | 0.574861 | 0.553604 | 0.596118 | 0.578273 | 0.565429 | 0.149968 | 0.600651 |
| $T\Delta\Delta S_{conf}$,N-D | 0.599168 | 0.575231 | 0.623105 | 0.604164 | 0.589203 | 0.19864 | 0.632708 |
| **Horizontal features** | **0.716636** | **0.715896** | **0.717375** | **0.717161** | **0.716156** | **0.433723** | **0.792609** |
| **Total features** | **0.729945** | **0.735305** | **0.724584** | **0.727746** | **0.731189** | **0.460337** | **0.81032** |

**Table S8. Sub-predictor statistics for Tyrosine (Y) subclass.** Values in red font indicate the largest statistic value in each feature group.

| Class Tyr | Accuracy | Sensitivity | Specificity | Precision | F1 | MCC | AUROC |
|---|---|---|---|---|---|---|---|
| ZIMJ680104 | 0.619481 | 0.602857 | 0.636104 | 0.623745 | 0.612989 | 0.239191 | 0.665827 |
| FASG760101 | 0.577662 | 0.616104 | 0.539221 | 0.572463 | 0.593164 | 0.155954 | 0.609484 |
| GRAR740103 | 0.615325 | 0.604675 | 0.625974 | 0.618149 | 0.611077 | 0.230897 | 0.67007 |
| RADA880106 | 0.593636 | 0.549091 | 0.638182 | 0.60318 | 0.574597 | 0.188191 | 0.637699 |
| **Vertical features** | **0.678052** | **0.643377** | **0.712727** | **0.691855** | **0.666293** | **0.357387** | **0.741669** |
| GUYH850101 | 0.681429 | 0.746753 | 0.616104 | 0.660855 | 0.701002 | 0.366175 | 0.74083 |
| MIYS990104 | 0.692078 | 0.741299 | 0.642857 | 0.675707 | 0.706663 | 0.386327 | 0.756548 |
| PRAM900102 | 0.554545 | 0.439221 | 0.66987 | 0.571021 | 0.496193 | 0.1122 | 0.577257 |
| PALJ810112 | 0.628701 | 0.658961 | 0.598442 | 0.621619 | 0.639436 | 0.258134 | 0.678237 |
| ROBB760105 | 0.648442 | 0.681039 | 0.615844 | 0.640043 | 0.659655 | 0.297698 | 0.705415 |
| PPIIPRO | 0.585065 | 0.414545 | 0.755584 | 0.629146 | 0.49953 | 0.181036 | 0.619905 |
| $\Delta G$,N | 0.607143 | 0.661039 | 0.553247 | 0.596803 | 0.627118 | 0.21569 | 0.659779 |
| $\Delta H_{ap}$,N | 0.528701 | 0.406494 | 0.650909 | 0.538099 | 0.462391 | 0.059241 | 0.529689 |
| $\Delta H_{pol}$,N | 0.574026 | 0.605195 | 0.542857 | 0.569747 | 0.586784 | 0.148431 | 0.600974 |
| $T\Delta S_{conf}$,N | 0.57 | 0.603377 | 0.536623 | 0.565604 | 0.583715 | 0.14041 | 0.595487 |
| $\Delta\Delta G$,N-D | 0.581688 | 0.592208 | 0.571169 | 0.580186 | 0.585936 | 0.163518 | 0.622842 |
| $\Delta\Delta H_{ap}$,N-D | 0.613247 | 0.672208 | 0.554286 | 0.601535 | 0.634806 | 0.228138 | 0.657273 |
| $\Delta\Delta H_{pol}$,N-D | 0.550649 | 0.560519 | 0.540779 | 0.549677 | 0.554867 | 0.101372 | 0.581004 |
| $T\Delta\Delta S_{conf}$,N-D | 0.591169 | 0.605714 | 0.576623 | 0.58918 | 0.597043 | 0.182583 | 0.630948 |
| **Horizontal features** | **0.694675** | **0.713247** | **0.676104** | **0.688193** | **0.700174** | **0.390005** | **0.76217** |
| **Total features** | **0.718442** | **0.717143** | **0.71974** | **0.719261** | **0.717993** | **0.43715** | **0.791034** |

**Table S9. Full PHOSforUS predictor performances calculated from X10 cross-validation.**

| Class S-P | Accuracy | Sensitivity | Specificity | Precision | F1 | MCC | AUROC |
|---|---|---|---|---|---|---|---|
| Vertical features | 0.752725 | 0.738784 | 0.766667 | 0.760029 | 0.749216 | 0.505698 | 0.835519 |
| Horizontal features | 0.78207 | 0.792733 | 0.771406 | 0.77631 | 0.784389 | 0.564335 | 0.870907 |
| Total features | 0.794589 | 0.800158 | 0.789021 | 0.791433 | 0.795736 | 0.589268 | 0.882882 |
| **Class S-nP** | **Accuracy** | **Sensitivity** | **Specificity** | **Precision** | **F1** | **MCC** | **AUROC** |
| Vertical features | 0.791125 | 0.7627 | 0.81955 | 0.808718 | 0.785009 | 0.583229 | 0.873915 |
| Horizontal features | 0.818325 | 0.82745 | 0.8092 | 0.81277 | 0.819977 | 0.636863 | 0.898609 |
| Total features | 0.8376 | 0.8432 | 0.832 | 0.833922 | 0.838486 | 0.675324 | 0.918708 |
| **Class T-P** | **Accuracy** | **Sensitivity** | **Specificity** | **Precision** | **F1** | **MCC** | **AUROC** |
| Vertical features | 0.703295 | 0.709742 | 0.696848 | 0.700794 | 0.705197 | 0.406673 | 0.779973 |
| Horizontal features | 0.723782 | 0.760172 | 0.687393 | 0.708504 | 0.733173 | 0.449166 | 0.80064 |
| Total features | 0.741404 | 0.767908 | 0.7149 | 0.729282 | 0.747969 | 0.483678 | 0.8199 |
| **Class T-nP** | **Accuracy** | **Sensitivity** | **Specificity** | **Precision** | **F1** | **MCC** | **AUROC** |
| Vertical features | 0.697782 | 0.686322 | 0.709242 | 0.702684 | 0.694134 | 0.39596 | 0.767477 |
| Horizontal features | 0.716636 | 0.715896 | 0.717375 | 0.717161 | 0.716156 | 0.433723 | 0.792609 |
| Total features | 0.729945 | 0.735305 | 0.724584 | 0.727746 | 0.731189 | 0.460337 | 0.81032 |
| **Class Y** | **Accuracy** | **Sensitivity** | **Specificity** | **Precision** | **F1** | **MCC** | **AUROC** |
| Vertical features | 0.678052 | 0.643377 | 0.712727 | 0.691855 | 0.666293 | 0.357387 | 0.741669 |
| Horizontal features | 0.694675 | 0.713247 | 0.676104 | 0.688193 | 0.700174 | 0.390005 | 0.76217 |
| Total features | 0.718442 | 0.717143 | 0.71974 | 0.719261 | 0.717993 | 0.43715 | 0.791034 |

**Table S10. Full comparative analysis data of PHOSforUS with current phosphorylation site predictors.**

| Class S-nP | Accuracy | Sensitivity | Specificity | Precision | F1 score | MCC | AUROC |
|---|---|---|---|---|---|---|---|
| PHOSforUS | 0.795 | 0.74 | 0.85 | 0.834396 | 0.782946 | 0.595424 | 0.8707 |
| Disphos | 0.717 | 0.544 | 0.89 | 0.833368 | 0.657429 | 0.463373 | 0.82301 |
| Musite | 0.669 | 0.448 | 0.89 | 0.804146 | 0.574517 | 0.377379 | 0.78346 |
| Netphos3.1 | 0.616 | 0.862 | 0.37 | 0.57799 | 0.691847 | 0.266571 | 0.71711 |
| Rfphos | 0.637 | 0.372 | 0.902 | 0.791095 | 0.504715 | 0.322944 | 0.74387 |
| PhosPred-RF | 0.772 | 0.654 | 0.89 | 0.857754 | 0.740594 | 0.561035 | 0.81251 |
| PhosphoSVM | 0.656 | 0.366 | 0.946 | 0.873751 | 0.515124 | 0.383791 | 0.81356 |
| **Class T-nP** | **Accuracy** | **Sensitivity** | **Specificity** | **Precision** | **F1 score** | **MCC** | **AUROC** |
| PHOSforUS | 0.687 | 0.64 | 0.734 | 0.706602 | 0.671354 | 0.375909 | 0.74322 |
| Disphos | 0.578 | 0.37 | 0.786 | 0.632685 | 0.466229 | 0.171279 | 0.62811 |
| Musite | 0.599 | 0.366 | 0.832 | 0.684903 | 0.475075 | 0.223637 | 0.67413 |
| Netphos3.1 | 0.531 | 0.598 | 0.464 | 0.528168 | 0.560629 | 0.062365 | 0.53124 |
| Rfphos | 0.61 | 0.372 | 0.848 | 0.711223 | 0.486631 | 0.25072 | 0.67351 |
| PhosPred-RF | 0.666 | 0.578 | 0.754 | 0.701385 | 0.633465 | 0.337389 | 0.71469 |
| PhosphoSVM | 0.603 | 0.288 | 0.918 | 0.779423 | 0.419856 | 0.265534 | 0.72002 |
| **Class Y** | **Accuracy** | **Sensitivity** | **Specificity** | **Precision** | **F1 score** | **MCC** | **AUROC** |
| PHOSforUS | 0.663 | 0.588 | 0.738 | 0.693753 | 0.634787 | 0.331058 | 0.72352 |
| Disphos | 0.595 | 0.412 | 0.778 | 0.653431 | 0.504334 | 0.2056 | 0.65703 |
| Musite | 0.6 | 0.578 | 0.622 | 0.606359 | 0.590998 | 0.200748 | 0.65107 |
| Netphos3.1 | 0.603 | 0.55 | 0.656 | 0.616933 | 0.580175 | 0.208103 | 0.62247 |
| Rfphos | 0.594 | 0.476 | 0.712 | 0.62333 | 0.539352 | 0.193675 | 0.63655 |
| PhosPred-RF | 0.62 | 0.744 | 0.496 | 0.596424 | 0.662059 | 0.247556 | 0.68372 |
| PhosphoSVM | 0.619 | 0.686 | 0.552 | 0.604983 | 0.642854 | 0.240269 | 0.67874 |
| **Class S-P** | **Accuracy** | **Sensitivity** | **Specificity** | **Precision** | **F1 score** | **MCC** | **AUROC** |
| PHOSforUS | 0.763 | 0.72 | 0.806 | 0.787563 | 0.752005 | 0.528191 | 0.84546 |
| Disphos | 0.69 | 0.692 | 0.688 | 0.691998 | 0.691489 | 0.380551 | 0.75849 |
| Musite | 0.631 | 0.868 | 0.394 | 0.591041 | 0.702042 | 0.297662 | 0.71465 |
| Netphos3.1 | 0.532 | 0.972 | 0.092 | 0.517175 | 0.67509 | 0.130089 | 0.63346 |
| Rfphos | 0.608 | 0.836 | 0.38 | 0.57441 | 0.680826 | 0.242584 | 0.67445 |
| PhosPred-RF | 0.66 | 0.95 | 0.37 | 0.602202 | 0.73677 | 0.39284 | 0.73841 |
| PhosphoSVM | 0.553 | 0.98 | 0.126 | 0.528897 | 0.686878 | 0.201449 | 0.70333 |
| **Class T-P** | **Accuracy** | **Sensitivity** | **Specificity** | **Precision** | **F1 score** | **MCC** | **AUROC** |
| PHOSforUS | 0.69 | 0.666 | 0.714 | 0.700031 | 0.682284 | 0.380759 | 0.76799 |
| Disphos | 0.592 | 0.762 | 0.422 | 0.568356 | 0.650632 | 0.197605 | 0.66303 |
| Musite | 0.597 | 0.838 | 0.356 | 0.565398 | 0.675205 | 0.221692 | 0.6354 |
| Netphos3.1 | 0.52 | 0.95 | 0.09 | 0.510748 | 0.664287 | 0.079128 | 0.59619 |
| Rfphos | 0.583 | 0.87 | 0.296 | 0.552847 | 0.675764 | 0.204821 | 0.6442 |
| PhosPred-RF | 0.592 | 0.956 | 0.228 | 0.553552 | 0.700941 | 0.269366 | 0.68277 |
| PhosphoSVM | 0.59 | 0.956 | 0.224 | 0.551987 | 0.699722 | 0.26752 | 0.69151 |
